# Single-cell transcriptomics and machine learning reveal RNF144B and C5AR1 as immune-related biomarkers and therapeutic targets in myocardial infarction

**DOI:** 10.1101/2025.09.03.673891

**Authors:** Yuxin Hu, Lu Chen, Jianmei Sha, Caihong Shao, Junli Gao, Jianhua Yao

**Affiliations:** Cardiac Regeneration and Ageing Lab, Institute of Geriatrics (Shanghai University), Affiliated Nantong Hospital of Shanghai University (The Sixth People’s Hospital of Nantong) and School of Life Science, Shanghai University, Nantong 226011, China; Institute of Cardiovascular Sciences, Shanghai Engineering Research Center of Organ Repair, Joint International Research Laboratory of Biomaterials and Biotechnology in Organ Repair (Ministry of Education), School of Life Science, Shanghai University, Shanghai 200444, China; Shanghai University Hospital, Shanghai University, Shanghai 200444, China; Department of Internal Emergency Medicine, Shanghai East Hospital, School of Medicine, Tongji University, Shanghai, 200120, China; Department of Cardiology, Tenth People’s Hospital, School of Medicine, Tongji University, Shanghai, 200090, China

**Author notes:** These authors contributed equally to this work (Yuxin Hu, Lu Chen, Jianmei Sha). Correspondence to: Dr. Junli Gao, Dr. Jianhua Yao.

## Abstract

**Background:** Myocardial infarction (MI) is a life-threatening cardiovascular disease characterized by high morbidity and mortality. Although advances in clinical management have improved patient outcomes, early diagnosis and effective immunomodulatory therapies remain limited.

**Methods:** In this study, we integrated multiple transcriptomic datasets and applied machine learning approaches, including LASSO regression, to identify a robust 13 key genes significantly associated with MI. Gene Set Enrichment Analysis (GSEA) and Gene Set Variation Analysis (GSVA) were subsequently performed to explore their potential biological functions. The immunological relevance of these genes was evaluated by analyzing their correlations with inflammation-related genes and those involved in immune cell migration. In addition, transcription factor (Johnson, Law et al.) and microRNA (miRNA) regulatory networks were constructed to elucidate upstream regulatory mechanisms. The expression levels of the 13 key genes were validated in MI mouse model. Furthermore, molecular docking was performed to identify candidate small molecules targeting core genes.

**Results:** Among 11 cardiac cell populations identified, myeloid cells contributed most prominently to MI pathogenesis. A robust 13-gene predictive signature was established, with RNF144B and C5AR1 showing strong associations with immune modulation and disease severity. GSEA and GSVA further revealed that RNF144B was enriched in the neutrophil degranulation pathway, while C5AR1 was associated with the complement cascade. Correlation analysis demonstrated a significant positive relationship of RNF144B and C5AR1 with immunological roles. Both genes were also positively correlated with classical MI marker genes SERPINE1 and RUNX1. TF-gene and miRNA–mRNA regulatory networks supported the post-transcriptional regulation of these genes. In the MI mouse model, expression of the 13 genes was consistent with the risk-prediction model. Molecular docking identified CCX168 as a promising small-molecule candidate targeting RNF144B and C5AR1.

**Conclusion:** This study reveals immune-related transcriptional signatures and signaling pathways that drive MI progression. The identified 13-gene signature, particularly RNF144B and C5AR1, holds promise as a diagnostic biomarker and therapeutic target, providing new insights for immunomodulatory and precision medicine strategies in MI. Keywords: Immune microenvironment, myeloid cells, inflammation, RNF144B, MI, immune-targeted therapy

**Highlights:**

- A 13-gene signature was identified to predict MI risk
- RNF144B and C5AR1 are key immune regulators in MI progression
- GSEA and GSVA reveal immune-related pathways of RNF144B and C5AR1
- CCX168 is identified as a potential small-molecule therapy for MI
- Mouse model validates gene expression consistent with the prediction model

## 1. Introduction

MI remains a leading cause of morbidity and mortality worldwide(Kathiresan and Srivastava 2012, Salari, Morddarvanjoghi et al. 2023). According to the World Health Organization, ischemic heart diseases, account for approximately 16% of all global deaths, underscoring the critical public health burden posed by this condition (Yokota, McCourt et al. 2020, Huynh, Hoffmann et al. 2024). Acute MI is commonly precipitated by sudden coronary artery thrombosis and occlusion, leading to ischemic injury, subsequent scar formation, and potential progression to heart failure(Qian 2016, Reboll, Klede et al. 2022, Virani 2023). Despite advancements in treatment, post-MI mortality rates remain significantly high, with estimates ranging from 7% to 20% within the first year following the MI event(Yokota, McCourt et al. 2020). This is associated with adverse cardiac remodeling, including ventricular dilation and fibrosis, which impairs cardiac function(Nguyen, Canseco et al. 2020, Kramann 2022).

Currently, the diagnosis of MI relies heavily on a combination of clinical symptoms, electrocardiographic changes, and the measurement of cardiac biomarkers such as troponins(McCarthy, Vaduganathan and Januzzi 2018). While these approaches are effective in many cases, they often lack the sensitivity and specificity required for early-stage detection or for distinguishing MI from other cardiovascular and systemic conditions(Schlotter, Huber et al. 2024). This diagnostic limitation hampers timely therapeutic interventions, which are critical for reducing myocardial damage and improving patient outcomes. Timely coronary reperfusion through interventions like stenting and coronary artery bypass grafting is standard in clinical practice to limit infraction extent. However, not all patients are suitable candidates for these therapies(Das, Goldstone et al. 2019). Therapeutic manipulation of the repair process, driven by robust tissue inflammation and its subsequent resolution, remains challenging(Prabhu and Frangogiannis 2016). Therefore, there is a pressing need to identify novel biomarkers that can enable early and accurate detection of MI and provide deeper insights into its underlying mechanisms.

In recent years, the integration of high-throughput sequencing technologies and computational analysis has revolutionized cardiovascular research(Kuppe, Ramirez Flores et al. 2022). Transcriptomic profiling has facilitated the identification of gene expression changes associated with disease states, while scRNA-seq offers unprecedented resolution to explore cellular heterogeneity and dynamics within complex tissues such as the heart (Ruiz-Villalba, Romero et al. 2020, Feng, Li et al. 2024, Wang and Dou 2024). Moreover, the application of machine learning algorithms has emerged as a powerful strategy to identify robust biomarkers and construct predictive models with high diagnostic accuracy (Martini 2023, Vergallo and Patrono 2023). These approaches hold great promise in bridging the gap between molecular research and clinical application(Kramann 2022).

In this study, we utilized scRNA-seq technology to compare cell subtypes between infracted and remote zones of MI-affected hearts, performing cell-type annotation and screening for genes closely associated with disease progression. To pinpoint key regulatory genes, we employed the LASSO regression, yielding a 13-gene predictive signature with high accuracy and stability. Among the identified genes, **RNF144B** emerged as a particularly promising biomarker due to its strong correlation with MI development. The correlations between these key genes and immune cells were analyzed via the single-sample gene set enrichment analysis (ssGSEA) algorithm. Additionally, the roles key genes in disease progression were investigated through gene set enrichment and transcription factor regulatory network analyses. Finally, by constructing a gene coexpression network, we validated the associations between the 13 key genes and the well-characterized MI-related genes RUNX1(McCarroll, He et al. 2018, Riddell, McBride et al. 2020, Amrute, Luo et al. 2024) and SERPINE1(Morange, Saut et al. 2007, Pu, Bao et al. 2022), thereby offering deeper insights into the pathogenesis of myocardial infarction.

Collectively, our study analyzed single-cell data from MI patients, revealing crucial cell types, key genes, immune infiltration patterns, and significant regulatory mechanisms within signaling pathways during disease development. Our integrated approach not only highlights RNF144B, C5AR1 and 11 other key genes as potential diagnostic and therapeutic targets but also provides valuable insights into the cellular and molecular mechanisms underlying MI. These findings offer a foundation for developing more effective strategies for early detection, risk assessment, and personalized treatment of myocardial infarction.

## 2 Methods

### 2.1 Data acquisition

The data used in this study were obtained from the Gene Expression Omnibus (GEO) database (https://www.ncbi.nlm.nih.gov/geo/) and Zenodo repository. The single-cell data file **Zenodo data** was used for single-cell analysis with 31 samples and 23 patients(Kuppe, Ramirez Flores et al. 2022). The dataset GSE214611 samples were collected from MI mouse model(Calcagno, Taghdiri et al. 2022). The GSE216211 dataset was annotated by the GPL24247 platform(Xu, Jiang et al. 2023).

### 2.2 Single-cell analysis

Cells expressing fewer than 200 or more than 10,000 genes were excluded to remove noncellular debris and potential cell aggregates. The data were log-normalized, and highly variable genes were identified via the FindVariableFeatures function. Subsequent analyses, including data scaling and principal component analysis (PCA), were conducted via the ScaleData and RunPCA functions, respectively. Cell clustering was performed with the FindNeighbors and FindClusters functions, and uniform manifold approximation and projection (UMAP) was used for cluster visualization, implemented in the Seurat R package as per the official vignettes (https://satijalab.org/seurat/articles/get_started.html). Cluster annotation was performed on the basis of known cell markers.

### 2.3 Immuno infiltration analysis

ssGSEA is extensively used to assess the types of immune cells within microenvironments. This approach identifies 22 immune cell phenotypes, encompassing T cells, B cells, and NK cells, among others. In this study, the ssGSEA algorithm was applied to quantify immune cells within the expression profile, thereby estimating the relative abundances of **22** different types of infiltrating immune cells.

### 2.4 Construction of the prediction model

To select marker genes of key cells, we used the GSE214611 dataset as the training set, while the GSE216211 dataset was used as the validation set. The prediction-related models were constructed using machine learning LASSO regression. Following the integration of the expression value for each gene, a risk score formula was established for each individual and weighted according to the regression coefficient estimated through LASSO regression analysis. The risk score formula was applied to calculate the score for each patient. The precision of the model predictions was assessed by a receiver operating characteristic (ROC) curve.

### 2.5 Contribution of different cell subpopulations to DFU

To evaluate the contributions of distinct cell subsets to disease development, both changes in cell abundance and gene expression profiles were examined. Initially, marker genes for each subset were identified through differential expression analysis. This involved selecting the top 100 genes with the highest expression in the control versus disease groups, which served as representative genes for each condition. Following this, the expression differences and relative proportions of these genes within each cell type were quantified. Finally, the contribution of each cell subset to the disease was calculated using the square root of the product of fold change and proportion (FC × PctProp).

### 2.6 Gene set enrichment analysis (GSEA)

Patients were categorized into high and low expression groups based on their gene expression levels. To further explore differences in signaling pathways between these groups, gene set enrichment analysis (GSEA) was performed. The reference gene sets were sourced from the Molecular Signatures Database (MsigDB), specifically focusing on annotated subtype-related pathways. Differential expression analysis was applied to compare pathway activity across subtypes. Significantly enriched gene sets were identified using consistency scores with an adjusted p-value threshold of < 0.05. GSEA is commonly utilized in research that integrates disease classification with underlying biological mechanisms.

### 2.7 Regulatory network analysis of key genes

The prediction of transcription factors was performed using the R package ‘RcisTarget,’ which bases its calculations entirely on motif data. The normalized enrichment score (NES) assigned to each motif depends on the total number of motifs present in the database. Beyond the motifs originally annotated in the dataset, additional annotations were inferred through analyses of motif similarity and gene sequence information. To assess motif enrichment within a gene set, the area under the curve (AUC) was computed for each motif-to-motif set comparison, utilizing recovery curve analysis that evaluates the gene set relative to the motif rankings. Subsequently, the NES for each motif was calculated from the distribution of AUC values across all motifs in the gene set. For the gene motif ranking database, this study used RcisTarget.hg19.motifDBs.cisbpOnly.500bp.

### 2.8 Gene set variation analysis (GSVA)

Gene set variation analysis (GSVA) is a nonparametric and unsupervised method designed to assess the enrichment of predefined gene sets within transcriptomic datasets. By aggregating gene-level changes into pathway-level shifts, GSVA provides insight into the functional biology of samples through comprehensive scoring of targeted gene sets. In this study, gene sets were obtained from the Molecular Signatures Database (MSigDB, version 7.0), and GSVA was employed to calculate enrichment scores for each set with precision. This methodology facilitated the evaluation of potential differences in biological processes across distinct samples.

### 2.9 Statistical analysis

All statistical analyses were performed using R software (version 4.3). The dataset underwent thorough preprocessing steps, including imputation of missing data and logarithmic transformation, to enhance comparability across groups and improve analytical accuracy. For comparisons between two groups, the Wilcoxon rank-sum test was applied, appropriate for data not following a normal distribution, with statistical significance defined at p < 0.05. Furthermore, to develop a reliable predictive model, feature selection was conducted using the LASSO regression technique. The model’s diagnostic capability and stability were evaluated by receiver operating characteristic (ROC) curves and the corresponding area under the curve (AUC) metrics.

### 2.10 Animal studies

For this study, 8-week-old male C57BL/6J mice were used. The anesthesia agent used in the animal experiments was ketamine hydrochloride. All animal procedures were performed in accordance with protocols approved by Shanghai University Committee on Scientific Ethics.

### 2.11 Expression Validation of Key Genes via Quantitative PCR

To validate the expression of key genes identified through bioinformatics analysis between myocytes and normal tissue, transcriptomes from 4 Sham mice and 4 MI mice were analyzed. Total RNA was extracted from heart tissues via TRIzol reagent (Invitrogen, USA), followed by reverse transcription into cDNA via the PrimeScript RT Master Mix (Takara, Japan). Quantitative PCR (qPCR) was performed via a SYBR Premix Ex Taq II kit (Takara, Japan). Gene expression levels were normalized to that of GAPDH, which was used as the internal control.

## 3 Results

### 3.1 scRNA-seq analysis identify contribution of different cell subpopulations to MI

To identify novel predictive and therapeutic factors associated with MI, we conducted data mining using high quality scRNA-seq datasets. The primary scRNA-seq datasets were obtained from the Zenodo data repository (DOI: 10.5281/zenodo.6578047), comprising 31 samples from 23 MI patients. The dataset includes three distinct regions: infarct zone (IZ), border zone (BZ), and remote zone (RZ). In addition, fibrotic zone (FZ) samples-typically representing fibrotic tissue-were collected 30-200 days post-MI.

The dataset underwent standard preprocessing steps, including normalization, scaling, and principal component analysis (PCA), followed by batch correction using Harmony (Fig. S1d, e). Dimensionality reduction and clustering were performed using UMAP, which identified 11 distinct cell subpopulations (Fig. 1b). Based on the expression of known marker genes, the clusters were assigned to the following cell types: cardiac muscle myoblasts, cardiac fibroblasts, cardiac endothelial cells, immature innate lymphoid cells, pericytes, lymphoid lineage–restricted progenitor cells, smooth muscle myoblasts, neuronal receptor cells, myeloid cells, epicardial adipocytes from the left ventricle, and others. A bar plot showing the expression levels of classical marker genes across these 11 cell types is presented in Fig. 2d.

**Fig. 1.**
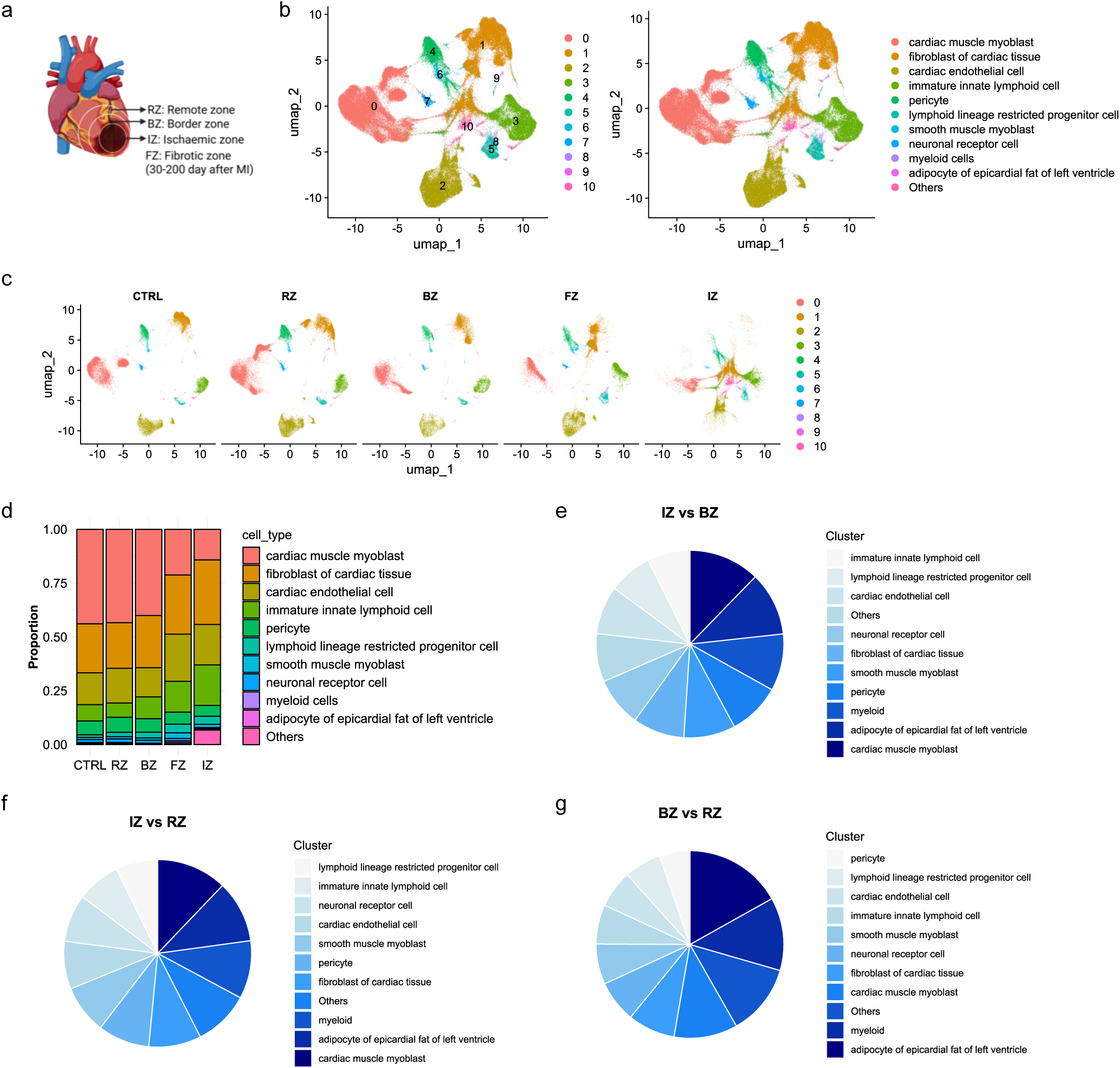
Single-cell sequencing of myocardial infarction (MI) tissues from different regions reveals a strong association between myeloid cells and the pathological progression of MI. a. Schematic illustration of regional partitioning for single-cell sequencing in MI tissues. RZ: remote zone; BZ: border zone; IZ: ischemic zone; FZ: fibrotic zone (30—200 days after MI). b. UMAP clustering analysis of single-cell sequencing data, identifying 11 distinct cell clusters. c. Distribution of cell clusters across different MI regions. d. Proportional changes in cell types across different MI regions. e-g. Contribution of different cell types to the pathological progression of MI.

**Fig. 2.**
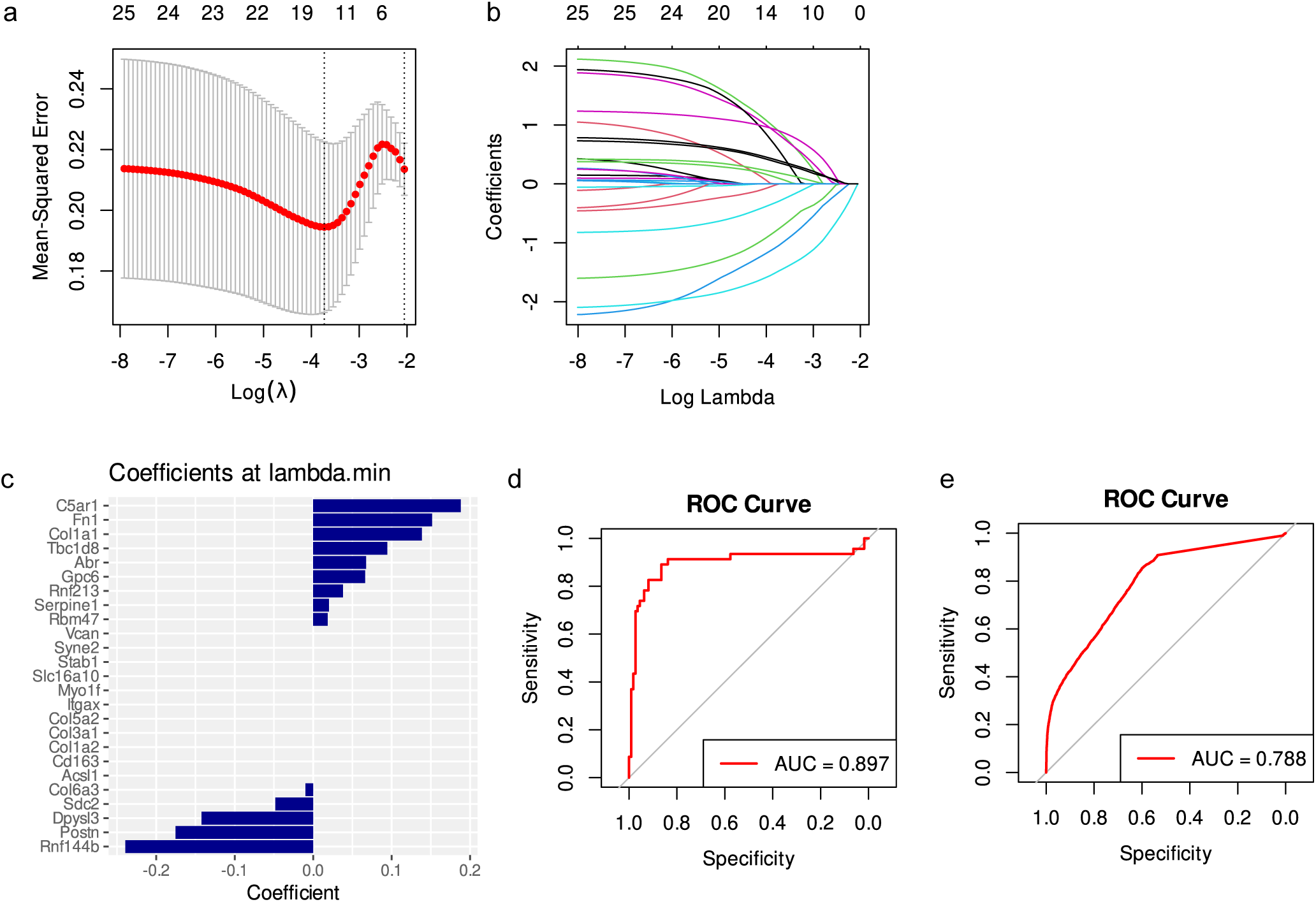
Construction of the 13-genes prediction model. a. Tenfold cross-validation was performed using the LASSO model to ascertain the minimum lambda value for parameter tuning. b. The distribution of LASSO coefficients and the gene combinations at the minimum lambda value were examined. c. The coefficients of the LASSO genes were analysed. d-e. Receiver operating characteristic curves of the model were generated.

To quantify the contribution of each cell type to disease pathology, we used the square root of (fold change × expression proportion) as a composite metric. Three pairwise comparisons were conducted: IZ vs. BZ, IZ vs. RZ, and BZ vs. RZ. To evaluate cell-type-specific contributions to disease progression, we first identified the top 100 highly expressed genes in both control and disease samples (total sample). We then calculated the differential expression and proportional expression of these genes within each cell subtype.

Among all subtypes, cardiac muscle myoblast, epicardial adipocytes of the left ventricle and myeloid cells exhibited the highest contribution scores across all three comparisons (Fig. 1e-g). Considering the limited abundance of epicardial adipocytes in cardiac tissue (constituting ∼504 cells at baseline) and their significant downregulation post-MI (reduction factor ∼0.5), we prioritized myeloid cells for subsequent analyses due to their greater cellular prevalence (∼701 cells at baseline) and marked upregulation following MI (induction factor ∼1.5). The IZ region represents the core area most affected by MI and reflects the most significant pathological alterations. Therefore, we selected IZ and RZ as the primary regions for further investigation.

We also explored the differences in metabolic pathway activity in myeloid cells between the IZ and RZ regions. The results revealed significant alterations in several pathways, including butanoate metabolism, beta-alanine metabolism, sulfur metabolism, fatty acid metabolism, ether lipid metabolism, and selenoamino acid metabolism, among others (Fig. S2d).

### 3.2 Construction of 13-gene prediction model of MI

To training the prediction model, we selected the GSE214611 dataset was used as the training set, while GSE216211 served as the validation set. Marker genes from 701 myeloid cells were identified and subjected to LASSO regression for feature selection. This process resulted in the identification of 13 genes characteristic of MI, which were subsequently used to construct a predictive model (Fig. 2a, b).

The model formula was defined as follows:

Risk score = –3.932573 + 0.958126 × FN1 – 1.091915 × POSTN + 0.118292 × RBM47 + 0.890296 × COL1A1 + 0.666463 × TBC1D8 + 0.198664 × RNF213 – 0.324249 × SDC2 + 0.414865 × ABR – 0.023176 × COL6A3 – 0.814206 × DPYSL3 – 1.516445 × RNF144B + 0.925670 × C5AR1 + 0.380113 × GPC6 (Fig. 2c).

The model exhibited strong diagnostic performance, with an area under the curve (AUC) of 0.897 in the training set (Fig. 2d). Validation using the external GSE216211 dataset confirmed the robustness of the model, achieving an AUC of 0.788 (Fig. 2e). The final 13 key genes included: FN1, POSTN, RBM47, COL1A1, TBC1D8, RNF213, SDC2, ABR, COL6A3, DPYSL3, RNF144B, C5AR1, and GPC6.

### 3.3 Immune cell infiltration in MI

As mentioned before, we found that immune cell myeloid strongly contribute to the pathological process of MI. To explore how immune system effects the MI progression, we analyzed immune cell infiltration patterns in the MI dataset.

Our results revealed the distribution of immune cell types across individual patients and uncovered the correlations among different immune cell populations (Fig. 3a, b). Notably, significant differences in the abundance of myeloid cells, including dendritic cells (DCs), M2 macrophages, and natural killer (NK) cells were observed between the two groups (Fig. 3c).

**Fig. 3.**
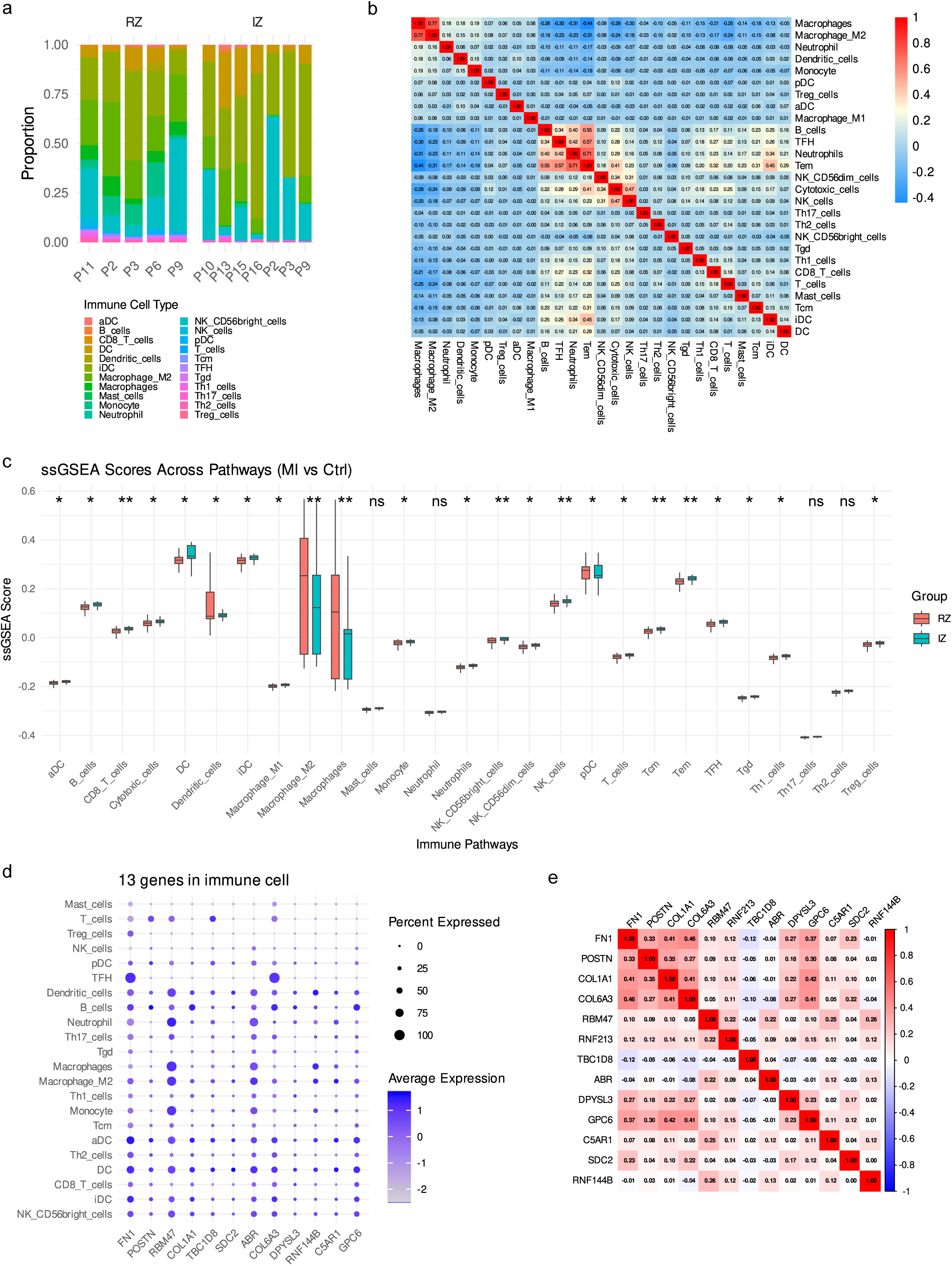
Immune cell infiltration analysis. **a**. Relative percentages of 22 immune cell subgroups.b. Pearson correlation coefficients for 22 types of immune cells, where blue signifies a negative correlation and red signifies a positive correlation. c. Variations in immune cell composition between the two samples. d. Correlations between key genes and immune cells. e. Correlation map of key gene expressions.

Furthermore, correlation analysis between the 13 key genes and immune cell types demonstrated strong associations. Specifically: FN1 and COL6A3 showed a strong positive correlation with T follicular helper (TFH) cells; POSTN and TBC1D8 exhibited a significant negative correlation with T cells; RBM47, ABR, and RNF144B were positively correlated with macrophages; COL1A1 and DPYSL3 showed positive correlations with B cells; SDC2, C5AR1, and GPC6 were positively correlated with dendritic cells (Fig. 3d). A strong inter-gene correlation was observed among the 13 key genes, suggesting potential co-regulation or shared biological functions (Fig. 3e).

These findings provide insight into the immunological landscape of MI and suggest potential pathways through which the identified key genes may contribute to disease pathogenesis via modulation of immune responses.

### 3.4 Pathway enrichment analyses identify immune-related key genes RNF144B and C5AR1 in MI

To elucidate the mechanisms through which the 13 key genes may influence MI progression, we first performed Gene Set Enrichment Analysis (GSEA). Notably, multiple genes—including C5AR1 and RNF144B—were significantly enriched in the B cell receptor (BCR) signaling complex, suggesting potential involvement in B cell–mediated immune responses (Fig. 4). Additionally, RNF144B was also enriched in PIP3 signaling in B lymphocytes, indicating a broader role in immune regulation. The BCR signaling pathway is directly interconnected with myeloid cells. Myeloid cells—such as dendritic cells and macrophages—can phagocytose pathogens, process antigens, and present antigenic peptides to B cells via MHC class II molecules, thereby promoting BCR-mediated antigen recognition, activation, and clonal expansion. Additionally, myeloid cells secrete cytokines such as IL-6, BAFF, and APRIL, which further enhance BCR signaling and promote the differentiation of B cells into plasma cells.

**Fig. 4.**
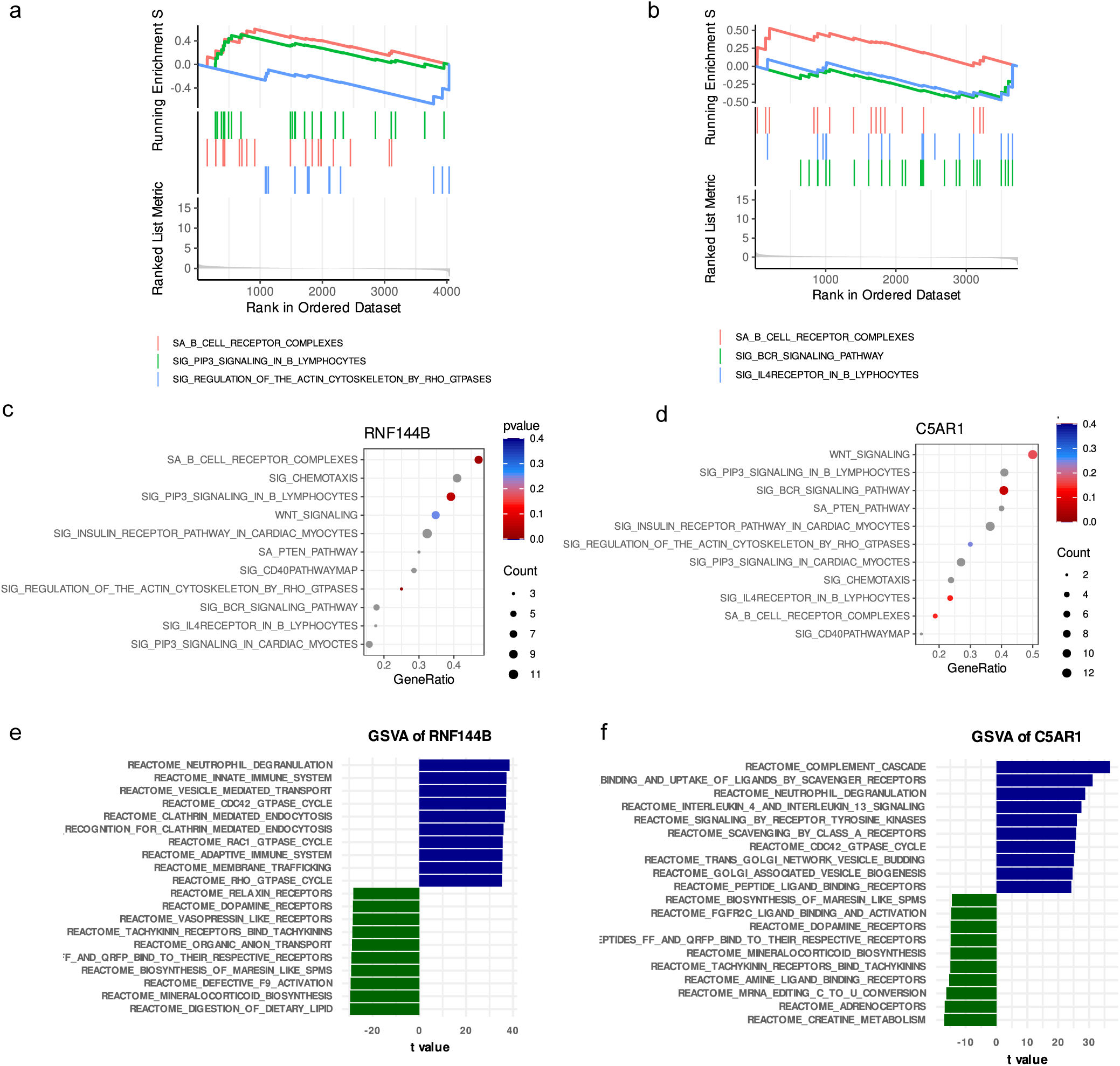
Among the 13-gene signature, RNF144B and C5AR1 show significant associations with immune-related pathways, such as the B-cell receptor signaling pathway. a-d. Gene Set Enrichment Analysis (GSEA) illustrating key genes involved in Kyoto Encyclopedia of Genes and Genomes (KEGG) signaling pathways, pathway regulation, and related genes. e-f. Gene Set Variation Analysis (GSVA). Blue indicates signaling pathways associated with high gene expression, while green represents pathways linked to low gene expression.

Complementary Gene Set Variation Analysis (GSVA) further supported immune pathway involvement. Specifically, RNF144B expression was positively associated with neutrophil degranulation, and C5AR1 was linked to the complement cascade (Fig. 4). These findings consistently highlight C5AR1 and RNF144B as key immune-related genes that may modulate the inflammatory microenvironment in MI.

### 3.5 RNF144B and C5AR1 are primarily involved in immune cells and pathways, and are positively correlated with MI-associated marker genes SERPINE1 and RUNX1

To further investigate the functional relevance of the key genes, we retrieved inflammation-related factors and immune cell migration–associated genes from the GeneCards database (https://www.genecards.org/). Correlation analyses were then performed between these functional gene sets and RNF144B and C5AR1 (Fig. 5, Fig. S6, Fig. S7).

**Fig. 5.**
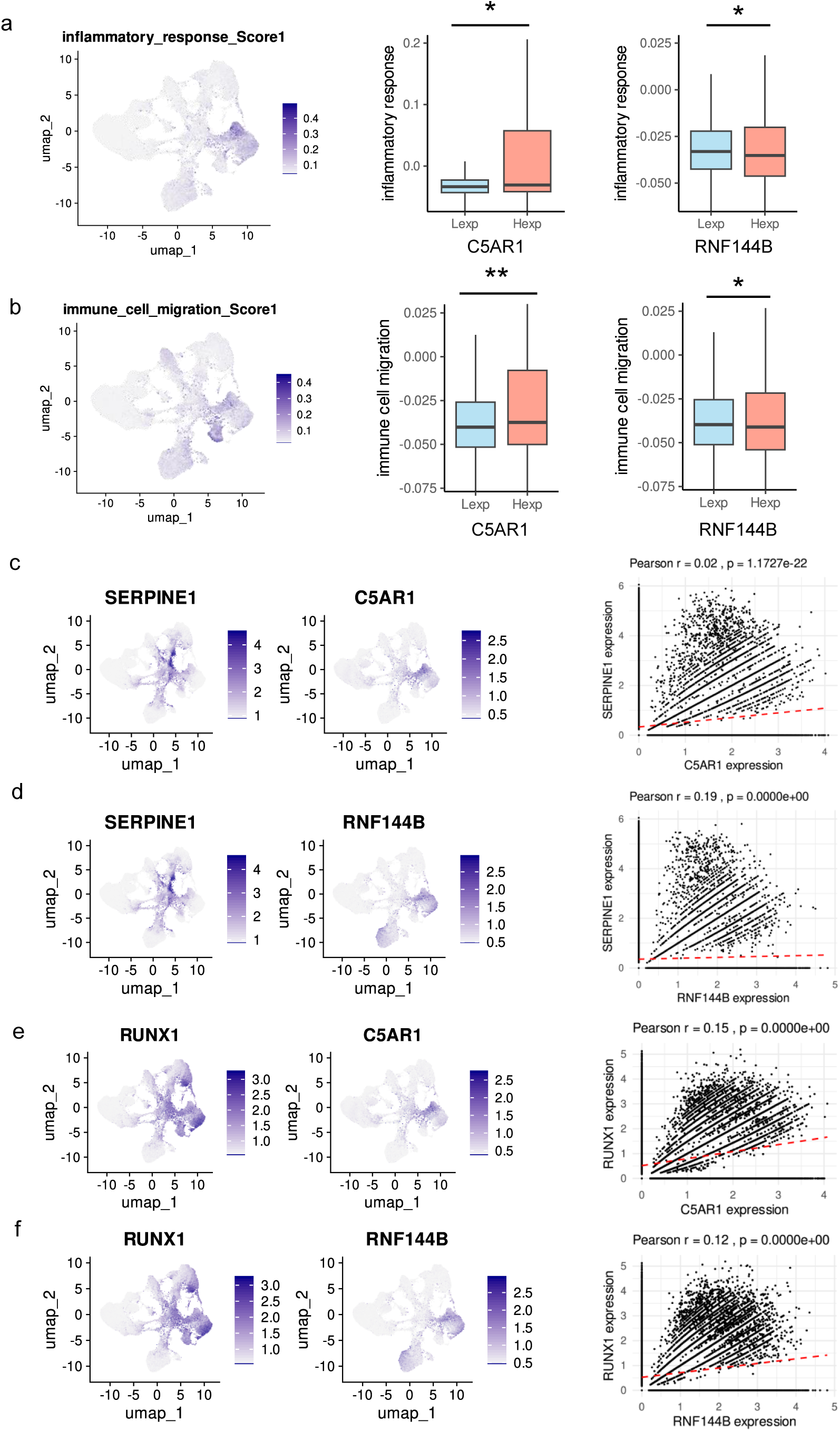
RNF144B and C5AR1 are associated with immune pathways and exhibit strong correlations with classical MI marker genes (SERPINE1 and RUNX1). a-b. Differences in the expression levels of C5AR1/RNF144B and inflammatory pathways, where red indicates high expression and blue represents low expression. c. co-expression analysis between C5AR1 and SERPINE1. d. co-expression analysis between RNF144B and SERPINE1. e. co-expression analysis between C5AR1 and RUNX1. f. co-expression analysis between RNF144B and RUNX1.

The results revealed that RNF144B expression was positively correlated with both inflammation-related factors and genes involved in immune cell migration. In contrast, C5AR1 exhibited a paradoxical pattern: its expression was downregulated in cells where inflammation and immune cell migration pathways were upregulated. This may reflect the cell type–specific functions of C5AR1—while it promotes pro-inflammatory responses in M1 macrophages and neutrophils, it exerts anti-inflammatory effects in M2 macrophages and myeloid-derived suppressor cells (MDSCs). Notably, both RNF144B and C5AR1 were positively correlated with two well-established MI marker genes, SERPINE1 and RUNX1, further supporting their potential involvement in MI-related immune regulation.

### 3.6 Regulatory network analysis of the 13 key genes

To explore the upstream regulatory mechanisms of the 13 key genes, we performed transcription factor (Johnson, Law et al.) enrichment analysis based on cumulative distribution curves. Motif annotation and enrichment analysis revealed several TF-binding motifs significantly associated with these genes. Among them, the motif taipale_tf_pairs_FOXO1_ELK3_RTMAACAGGAAGTN_CAP exhibited the highest normalized enrichment score (NES = 5.89), indicating a strong potential regulatory effect. All significantly enriched motifs and their corresponding transcription factors predicted to regulate the key genes are shown in Fig. 6a–c.

**Fig. 6.**
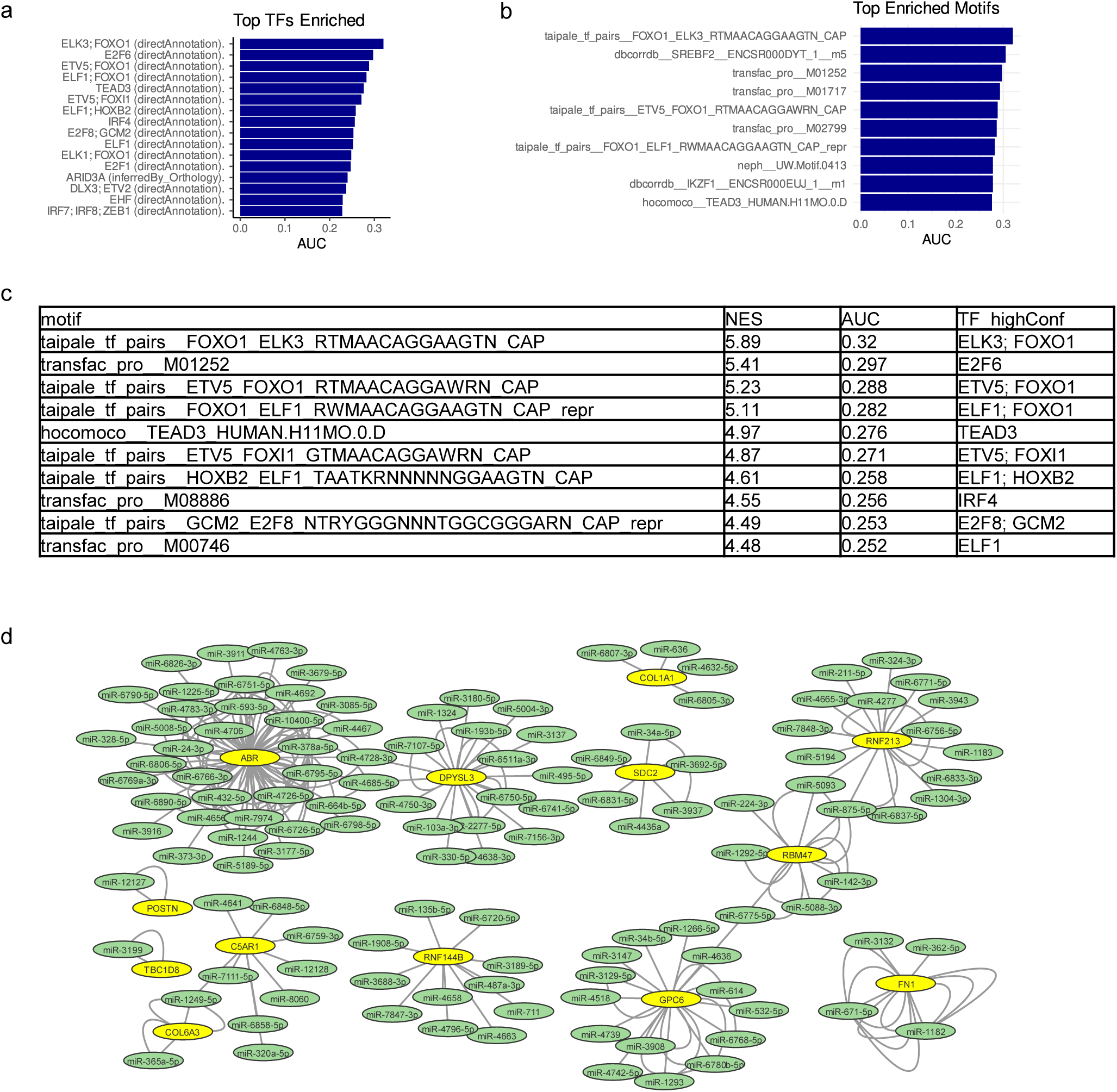
Key gene-related transcriptional regulation and miRNA networks. a. Transcriptional regulatory networks of key genes. b. All the enriched motifs, together with transcription factors associated with key genes. c-d. miRNA networks of key genes, with yellow representing mRNAs and green representing miRNAs.

In addition, a miRNA–mRNA regulatory network was constructed using the miRWalk database. A total of 224 predicted miRNA–mRNA interaction pairs involving the 13 key genes were identified and visualized with Cytoscape (Fig. 6d).

Together, these findings provide important insights into the transcriptional and post-transcriptional regulatory landscape of the key genes, highlighting their complex control in the context of myocardial infarction.

### 3.7 Validation of RNF144B and C5AR1 Expression and Molecular Docking Analysis in the MI Mouse Model

To validate the expression of the 13-gene predictive model, particularly RNF144B and C5AR1, qPCR was performed using MI mouse heart tissue (Fig. 7a–d). As predicted by the model, C5AR1 expression was significantly upregulated in the MI heart, whereas RNF144B expression was downregulated. The expression patterns of the remaining 11 genes were generally consistent with the model, with the exception of POSTN. This discrepancy may be attributed to the fact that the MI heart tissue was harvested 24 hours post-surgery, a time point at which fibrotic remodeling may not yet be fully established.

**Figure 7.**
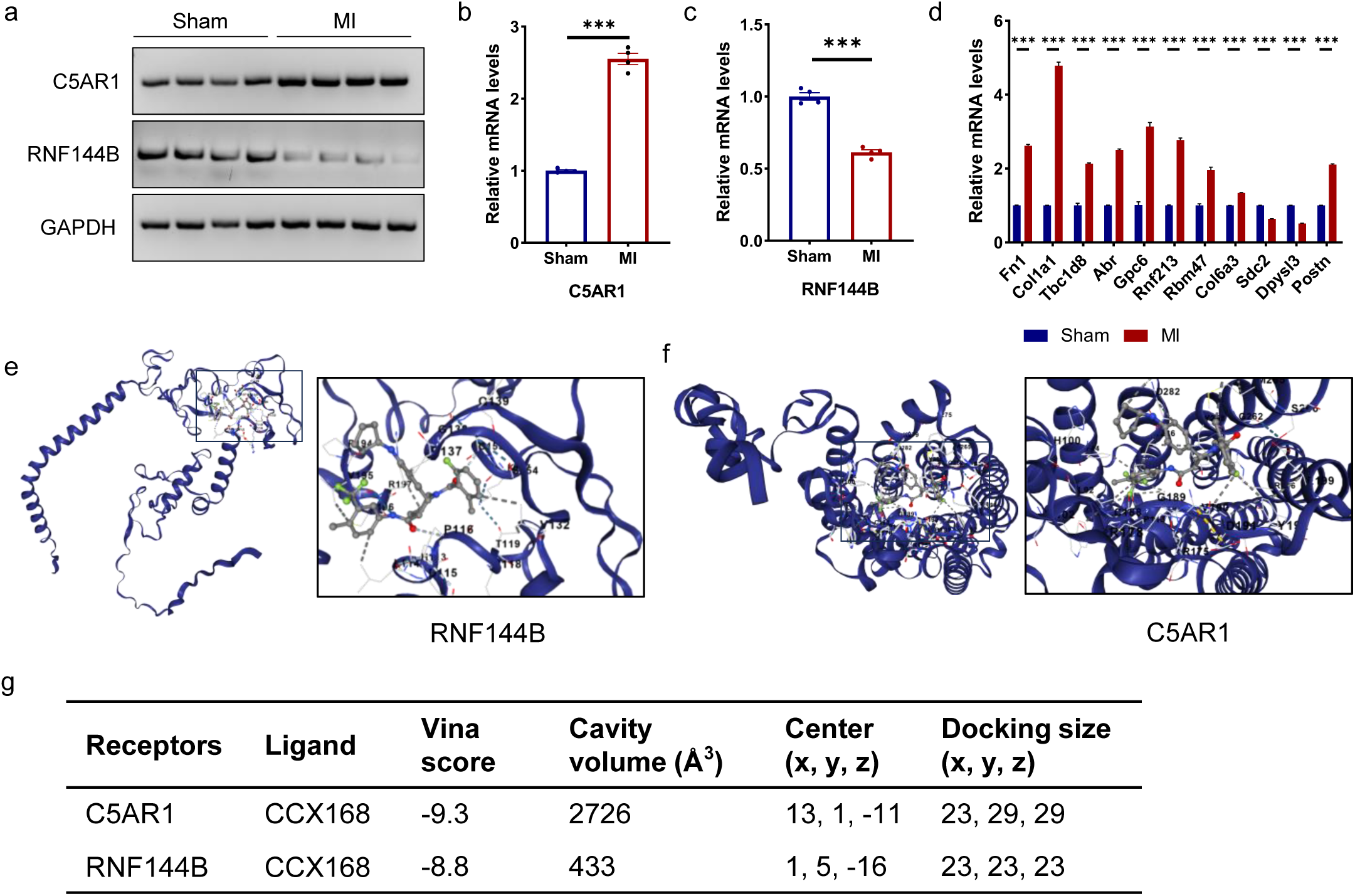
Validation of RNF144B and C5AR1 expression and docking analysis in the MI mouse model. a–c. Quantitative PCR (qPCR) validation of C5AR1 and RNF144B expression in the myocardial infarction (MI) mouse model. d. Expression levels of the remaining 11 signature genes in the MI model. e–f. Molecular docking of CCX168 with RNF144B and C5AR1. Protein structures are displayed as blue ribbons, and the docked ligands (CCX168) are shown in grey. g. Binding affinities between CCX168 and the core target proteins.

To further investigate potential interactions between the core target proteins (RNF144B and C5AR1) and candidate small-molecule compounds, molecular docking analysis was conducted. Candidate molecules were screened using the DIDB database, and CCX168 emerged as a promising ligand for both targets. CCX168 is the world’s first oral, selective inhibitor of the complement C5a receptor (C5aR), and has been approved for the treatment of two subtypes of ANCA-associated vasculitis (AAV): granulomatosis with polyangiitis (GPA) and microscopic polyangiitis (MPA). However, it has not yet been approved for use in cardiovascular or myocardial diseases.

Molecular docking analysis revealed that CCX168 exhibited strong binding affinities with both C5AR1 (Vina score: −9.3 kcal/mol) and RNF144B (Vina score: −8.8 kcal/mol) (Fig. 7e-g). A binding energy threshold of −5 kcal/mol was applied to indicate favorable interactions. The three-dimensional structures of RNF144B and C5AR1 were obtained from the AlphaFold Protein Structure Database. Docking simulations demonstrated stable and favorable binding of CCX168 to both targets. Notably, the binding site of CCX168 on RNF144B was located between amino acid residues 115–135, corresponding to the RING2 domain. This domain plays an essential role in stabilizing E2–Ub interactions and enhancing the catalytic ubiquitination activity of the RING1 domain. We hypothesize that CCX168 binding to the RING2 domain may enhance RNF144B’s E3 ligase activity by allosterically promoting RING1 function. The predicted binding pockets of RNF144B and C5AR1 are illustrated in Figure 7.

In summary, the expression of the 13-gene model was successfully validated in the MI mouse model, particularly RNF144B and C5AR1. Moreover, molecular docking analysis identified CCX168 as a potential candidate molecule for targeting these proteins, providing preliminary evidence for its possible therapeutic application in MI.

## 4. Discussion

MI arises from the occlusion of coronary arteries, depriving cardiac tissue of oxygen and leading to cell death, inflammation, and tissue remodeling. This intricate process involves the coordinated activity of multiple cell types, including cellular migration, proliferation, extracellular matrix deposition, and structural remodeling.

Our study revealed region-specific alterations in cellular composition within the heart tissue of MI patients. Notably, there was a marked increase in myeloid cells, immature innate lymphoid cells, and fibroblasts, accompanied by a reduction in cardiac myoblasts, pericytes, and neuronal receptor cells.

Among these, myeloid lineage cells—particularly macrophages, dendritic cells (DCs), granulocytes (neutrophils, eosinophils, basophils), monocytes, and mast cells—orchestrate indispensable functions across all stages of cardiac injury response, from initial inflammation to tissue repair and remodeling. In the setting of MI, macrophages show heightened inflammatory cytokine production and an altered M1/M2 polarization, skewed toward a proinflammatory M1 phenotype. Single-cell RNA sequencing (scRNA-seq) of mononuclear phagocytes in infarcted myocardium at day 11 post-ligation demonstrated the persistence of the four homeostatic macrophage clusters alongside four newly identified populations(Dick, Macklin et al. 2019). These MI-specific populations were enriched in pathways associated with inflammation, including TNF signaling, cell adhesion and migration, and hypoxia response(Peet, Ivetic et al. 2020).

Cardiac metabolism plays a central role in maintaining tissue homeostasis and orchestrating phenotypic changes in cells. The heart relies heavily on metabolic flexibility not only for ATP production to support contraction but also to drive biosynthetic processes essential for cell maintenance and regeneration. Disruptions in metabolic pathways during MI contribute to pathological remodeling, mediated by altered gene expression, redox imbalance, and impaired metabolic efficiency(Gibb and Hill 2018). Our findings align with previous studies showing that inefficient substrate utilization and diminished anabolic activity are proximate drivers of maladaptive cardiac remodeling.

To identify critical cellular contributors to MI, we screened for the top 100 highly expressed genes across cell types and assessed their expression and proportion changes. Myeloid cells emerged as the dominant contributors, underscoring their central role in post-infarction wound healing. We employed the LASSO regression algorithm, a robust machine learning tool for feature selection in high-dimensional datasets, to construct a predictive model based on myeloid-specific markers. Thirteen genes were identified and integrated into a diagnostic model that demonstrated strong performance and stability.

Among these genes, FN1, RBM47, COL1A1, TBC1D8, RNF213, ABR, C5AR1, and GPC6 were significantly downregulated in the infarct zone (IZ), while POSTN, SDC2, COL6A3, DPYSL3, and RNF144B were upregulated. Co-expression analysis highlighted strong correlations between FN1 and COL6A3, GPC6 and COL1A1, FN1 and DPYSL3, and FN1 and POSTN, suggesting that these gene pairs may cooperatively modulate inflammatory responses, hypoxia adaptation, and angiogenesis during MI progression.

Among these, RNF144B (Ring Finger Protein 144B) is a gene encoding a member of the RING-type E3 ubiquitin ligase family (Abad, Sandoz et al. 2024). These proteins are involved in the ubiquitination process, which tags proteins for degradation via the proteasome pathway. RNF144B is characterized by the presence of a RING finger domain, which facilitates its function in mediating protein-protein interactions and catalyzing the transfer of ubiquitin molecules. RNF144B has been implicated in various cellular processes, including apoptosis, immune response regulation(Li, Zhang et al. 2024), and tumorigenesis(Zhuang, Zhang et al. 2024). Emerging evidence suggests that RNF144B plays a role in modulating signaling pathways such as NF-κB and STAT, thereby influencing inflammation and immune homeostasis.

C5AR1 was the most significantly downregulated gene, indicating its pivotal role in MI pathogenesis. C5AR1, along with C5AR2, serves as a receptor for C5a, a central component of the complement cascade(Fattahi, Frydrych et al. 2018, Desai, Kumar et al. 2023, Liu, Chen et al. 2023). Beyond extracellular complement-mediated immunity, intracellular complement signaling—referred to as the “complosome”—has emerged as a key regulator of immunometabolism(Arbore, West et al. 2016, Nording, Baron et al. 2021). Macrophages can autonomously generate C5a, which activates C5AR1 on mitochondrial membranes, shifting energy production toward anaerobic glycolysis and reactive oxygen species generation, ultimately promoting IL-1β synthesis. Loss of macrophage-specific C5AR1 has been shown to alleviate cardiovascular disease and inflammation in various models, highlighting its potential as a therapeutic target in MI(Natarajan, Abbas et al. 2018, Niyonzima, Rahman et al. 2021). Therefore, C5AR1 is expected to become a potential therapeutic target for MI.

Periostin (POSTN), a 90-kDa matricellular protein, is minimally expressed in healthy adult myocardium but re-expressed following MI(Amrute, Luo et al. 2024). Although POSTN contributes to fibrosis and adverse remodeling, it has also been implicated in promoting cardiomyocyte proliferation and myocardial regeneration(Kanisicak, Khalil et al. 2016, Li, Lou et al. 2023). Administration of recombinant POSTN post-MI has been shown to enhance ventricular function, reduce infarct size, and increase angiogenesis, indicating a dual role in cardiac pathology and repair(Hudson and Porrello 2017).

Despite advances in medical management, current therapies for MI often fall short, partly due to an incomplete understanding of its underlying mechanisms. MI is characterized by excessive inflammation, ischemia, and hypoxia, all of which disrupt the immune microenvironment and cytokine homeostasis(Zhao, Williamson et al. 2024). Our immune infiltration analysis revealed substantial differences in immune cell subsets, including dendritic cells, macrophages, T follicular helper (TFH) cells, effector memory T (Tem) cells, NK cells, and monocytes between infarct (IZ) and remote zones (RZ). We also identified significant correlations between key genes and specific immune cell populations. T cells, in particular, serve as pivotal regulators of immune reactivity and tolerance(Fan, Tao et al. 2019, Jia, Chen et al. 2022). The MI microenvironment is marked by chronic inflammation and immune dysregulation, making targeted immunotherapy—particularly aimed at reducing inflammatory cell infiltration—a compelling strategy for future MI treatment.

GSEA further demonstrated that the key genes are predominantly enriched in the BCR and IL4R signaling pathways. The BCR pathway is central to antigen recognition, B cell activation, and humoral immunity IL4R signaling mediates type 2 immune responses, influencing B cell survival, class switching, and anti-inflammatory actions. Notably, IL4Rα signaling has shown cardioprotective effects in both mice and humans following ischemic injury(Alvarez-Argote, Almeida et al. 2024). These findings suggest that dysregulated BCR and IL4R signaling may contribute to impaired healing and persistent inflammation in MI.

This study has several limitations. The predictive model was constructed using the LASSO algorithm, which, while effective, may introduce bias if used in isolation. Incorporating additional machine learning methods (e.g., Random Forest, Support Vector Machine) could enhance model accuracy and generalizability. Moreover, although we identified key genes and pathways potentially involved in MI, our conclusions are primarily based on bioinformatics analyses and correlation-based inferences. Functional validation through in vitro and in vivo experiments is necessary to elucidate the precise roles and mechanisms of these genes in MI pathogenesis. Future work will focus on experimental validation and mechanistic exploration at the cellular and tissue levels.

### Clinical Relevance

Through single-cell RNA sequencing and transcriptomic analysis, a set of 13 genes were identified as pivotal contributors to the onset and progression of MI. Additionally, a prediction model was constructed using the LASSO regression algorithm. Finally, analysis of immune infiltration and signaling pathways revealed the potential mechanisms underlying MI development. The aforementioned findings offer innovative perspectives and avenues for forthcoming clinical investigations and therapeutic approaches.

## Glossary

Term / Abbreviation: Definition
MI (Myocardial Infarction): A condition commonly known as a heart attack, caused by the interruption of blood flow to the heart, resulting in tissue damage or necrosis.
scRNA-seq (Single-cell RNA sequencing): A high-throughput technique used to profile gene expression at the single-cell level, allowing analysis of cellular heterogeneity in complex tissues.
LASSO Regression (Least Absolute Shrinkage and Selection Operator): A machine learning algorithm used for feature selection and model construction by penalizing the absolute size of regression coefficients.
ssGSEA (Single-sample Gene Set Enrichment Analysis): A bioinformatics method for estimating the relative abundance of immune cell types within individual samples based on gene expression data.
GSEA (Gene Set Enrichment Analysis): A computational approach used to determine whether predefined gene sets show statistically significant differences between two biological states.
GSVA (Gene Set Variation Analysis): A non-parametric method that estimates variation of gene set enrichment across a sample population in an unsupervised manner.
RNF144B (Ring Finger Protein 144B): A gene encoding an E3 ubiquitin ligase involved in immune regulation and inflammation via NF-κB and STAT signaling pathways.
C5AR1 (Complement Component 5a Receptor 1): A G protein-coupled receptor for complement factor C5a, playing a critical role in complement activation, immune cell recruitment, and inflammation.
TF Network (Transcription Factor Network): A regulatory network representing the relationships between transcription factors and their target genes, influencing gene expression programs.
RUNX1 / SERPINE1: MI-associated genes known to regulate inflammation, fibrosis, and vascular remodeling; used here as reference markers for model validation.

## Statements and Declarations

### Ethics approval

We hereby confirm that the study reported in our manuscript has been conducted and reported in accordance with the ARRIVE (Animals in Research: Reporting In Vivo Experiments) guidelines (https://arriveguidelines.org). We hereby declare that all experimental procedures strictly followed international and institutional guidelines for animal research. The study protocol was approved by the Shanghai University Committee on Scientific Ethics (Approval No.: YS2024-195).

### Consent for publication

Not applicable.

### Availability of data and material

All data associated with this study are present in the paper or in the Supporting Information. The scRNA-seq data of keloids from the published paper are available in the GEO. Data, codes, and materials will be made available upon request.

### Author contributions

Gao and authors led the data analysis and manuscript writing. All the authors reviewed and approved the final version of the manuscript.

## Acknowledgments

This manuscript was supported by Dr. Xiao and Dr. Li.

## Authors’ information

1. Cardiac Regeneration and Aging Lab, Institute of Geriatrics (Shanghai University), Affiliated Nantong Hospital of Shanghai University (The Sixth People’s Hospital of Nantong) and School of Life Science, Shanghai University, Nantong 226011, China. 2. Institute of Cardiovascular Sciences, Shanghai Engineering Research Center of Organ Repair, Joint International Research Laboratory of Biomaterials and Biotechnology in Organ Repair (Ministry of Education), School of Life Science, Shanghai University, Shanghai 200444, China. 3. Shanghai University Hospital, Shanghai University, Shanghai 200444, China. 4. Department of Internal Emergency Medicine, Shanghai East Hospital, School of Medicine, Tongji University, Shanghai, 200120, China. 5Department of Cardiology, Tenth People’s Hospital, School of Medicine, Tongji University, Shanghai, 200090, China.

**No human studies were carried out by the authors for this article**

---

**Fig. S1 Single-cell analysis of MI**. a. Quality control for single-cell data involved the presentation of metrics such as cell count, gene count, and sequencing depth for each sample. b. The correlation between sequencing depth and mitochondrial content (left) and the correlation between sequencing depth and the number of genes (right); both are positively correlated. c. Gene feature variance plot showing significant differences between cells. d. Variance ranking chart for each PC. e. Visualisation of principal component analysis and distribution of PCs; colours stand for samples, and points stand for cells.

**Fig. S2. Analysis of the importance of cells and metabolic pathways.** a-c. The importance of 11 types of cells, from left to right, increases successively. d. Heatmap illustrating the correlation between two sample groups and metabolic pathways, with pink representing the control group and green representing the diabetic foot ulcer group

**Fig. S3 Expression of single cells.** (a, b) Expression of key genes in single cells

**Fig. S4 Gene set enrichment analysis of key genes.** a–j Gene Set Enrichment Analysis (GSEA) of selected genes showing immune and signaling pathway enrichment. GSEA identified that ABR was enriched in the B cell receptor complex and the insulin receptor signaling pathway in cardiac myocytes. RBM47, COL1A1, COL6A3, FN1, GPC6, and POSTN were enriched in the BCR signaling pathway. DPYSL3 and SDC2 were enriched in the IL-4 receptor signaling pathway in B lymphocytes, while TBC1D8 was enriched in the WNT signaling pathway.

**Fig. S5 Gene Set Variation Analysis (GSVA) based on expression levels of the 13 key genes.** Blue denotes the signaling pathways associated with high gene expression, and green indicates those linked to low gene expression. GSVA revealed that: ABR was associated with the NRAGE-mediated JNK signaling pathway;DPYSL3, with CRMPs in SEMA3A signaling;COL6A3, with collagen chain trimerization;COL1A1, with enhanced binding of GP1BA variant to VWF/collagen;FN1, with fibronectin matrix formation;GPC6 and SDC2, with the EXT2-deficiency pathway; POSTN, with collagen degradation;RBM47, with heme scavenging from plasma;TBC1D8, with vesicle-mediated transport;RNF213, with suppression of apoptosis.

**Fig. S6 Differences in the expression of key genes and inflammatory response scores.** Differences in the expression of key genes and cytokine scores, with red representing high expression and blue representing low expression.

**Fig. S7 Differences in the expression of key genes and inflammatory response scores.** Differences in the expression of key genes and cytokine scores, with red representing high expression and blue representing low expression.

**Fig. S8 Analysis of the coexpression of key genes and SERPINE1 in single cells.** The coexpression and correlation of each key gene with SERPINE1.

**Fig. S9 Analysis of the coexpression of key genes and RUNX1 in single cells.** The coexpression and correlation of each key gene with RUNX1.

## Reference

Abad, E., J. Sandoz, G. Romero, I. Zadra, J. Urgel-Solas, P. Borredat, S. Kourtis, L. Ortet, C. M. Martinez, D. Weghorn, S. Sdelci and A. Janic (2024). The TP53-activated E3 ligase RNF144B is a tumour suppressor that prevents genomic instability. Journal of experimental & clinical cancer research: CR 43(1): 127. 10.1186/s13046-024-03045-4.

Alvarez-Argote, S., V. A. Almeida, M. C. Knas, S. L. Buday, M. Patterson and C. C. O’Meara (2024). Global IL4Rα blockade exacerbates heart failure after an ischemic event in mice and humans. American Journal of Physiology-Heart and Circulatory Physiology 326(5): H1080–H1093. 10.1152/ajpheart.00010.2024

Amrute, J., X. Luo, V. Penna, S. Yang, T. Yamawaki, S. Hayat, A. Bredemeyer, I. Jung, F. Kadyrov, G. S. Heo, R. Venkatesan, S. Shi, A. Parvathaneni, A. Koenig, C. Kuppe, C. Baker, H. Luehmann, C. Jones, B. Kopecky, X. Zeng, T. Blekwehl, P. Ma, P. Lee, Y. Terada, A. Fu, M. Furtado, D. Kreisel, A. Kovacs, N. Stitziel, S. Jackson, C. M. Li, Y. J. Liu, N. Rosenthal, R. Kramann, B. Ason and K. Lavine (2024). Targeting Immune-Fibroblast Crosstalk in Myocardial Infarction and Cardiac Fibrosis. Circulation 150. 10.1161/circ.150.suppl_1.4140393

Amrute, J. M., X. Luo, V. Penna, S. Yang, T. Yamawaki, S. Hayat, A. Bredemeyer, I.-H. Jung, F. F. Kadyrov, G. S. Heo, R. Venkatesan, S. Y. Shi, A. Parvathaneni, A. L. Koenig, C. Kuppe, C. Baker, H. Luehmann, C. Jones, B. Kopecky, X. Zeng, T. Bleckwehl, P. Ma, P. Lee, Y. Terada, A. Fu, M. Furtado, D. Kreisel, A. Kovacs, N. O. Stitziel, S. Jackson, C.-M. Li, Y. Liu, N. A. Rosenthal, R. Kramann, B. Ason and K. J. Lavine (2024). Targeting immune-fibroblast cell communication in heart failure. Nature 635(8038): 423–433. 10.1038/s41586-024-08008-5

Arbore, G., E. E. West, R. Spolski, A. A. B. Robertson, A. Klos, C. Rheinheimer, P. Dutow, T. M. Woodruff, Z. X. Yu, L. A. O’Neill, R. C. Coll, A. Sher, W. J. Leonard, J. Kohl, P. Monk, M. A. Cooper, M. Arno, B. Afzali, H. J. Lachmann, A. P. Cope, K. D. Mayer-Barber and C. Kemper (2016). T helper 1 immunity requires complement-driven NLRP3 inflammasome activity in CD4_ T cells. Science (New York, N Y) 352(6292): aad1210. 10.1126/science.aad1210

Calcagno, D. M., N. Taghdiri, V. K. Ninh, J. M. Mesfin, A. Toomu, R. Sehgal, J. Lee, Y. Liang, J. M. Duran, E. Adler, K. L. Christman, K. Zhang, F. Sheikh, Z. Fu and K. R. King (2022). Single-cell and spatial transcriptomics of the infarcted heart define the dynamic onset of the border zone in response to mechanical destabilization. Nat Cardiovasc Res 1(11): 1039–1055. 10.1038/s44161-022-00160-3

Das, S., A. B. Goldstone, H. J. Wang, J. Farry, G. D’Amato, M. J. Paulsen, A. Eskandari, C. E. Hironaka, R. Phansalkar, B. Sharma, S. Rhee, E. A. Shamskhou, D. Agalliu, V. D. Perez, Y. J. Woo and K. Red-Horse (2019). A Unique Collateral Artery Development Program Promotes Neonatal Heart Regeneration. Cell 176(5): 1128-+. 10.1016/j.cell.2018.12.023

Desai, J. V., D. Kumar, T. Freiwald, D. Chauss, M. D. Johnson, M. S. Abers, J. M. Steinbrink, J. R. Perfect, B. Alexander, V. Matzaraki, B. D. Snarr, M. A. Zarakas, V. Oikonomou, L. M. Silva, R. Shivarathri, E. Beltran, L. N. Demontel, L. Wang, J. K. Lim, D. Launder, H. R. Conti, M. Swamydas, M. T. McClain, N. M. Moutsopoulos, M. Kazemian, M. G. Netea, V. Kumar, J. Kohl, C. Kemper, B. Afzali and M. S. Lionakis (2023). C5a-licensed phagocytes drive sterilizing immunity during systemic fungal infection. Cell 186(13): 2802–2822.e2822. 10.1016/j.cell.2023.04.031

Dick, S. A., J. A. Macklin, S. Nejat, A. Momen, X. Clemente-Casares, M. G. Althagafi, J. M. Chen, C. Kantores, S. Hosseinzadeh, L. Aronoff, A. Wong, R. Zaman, I. Barbu, R. Besla, K. J. Lavine, B. Razani, F. Ginhoux, M. Husain, M. I. Cybulsky, C. S. Robbins and S. Epelman (2019). Self-renewing resident cardiac macrophages limit adverse remodeling following myocardial infarction. Nature Immunology 20(1): 29-+. 10.1038/s41590-018-0272-2

Fan, Q., R. Tao, H. Zhang, H. Y. Xie, L. Lu, T. Wang, M. Su, J. Hu, Q. Zhang, Q. J. Chen, Y. Iwakura, W. F. Shen, R. Y. Zhang and X. X. Yan (2019). Dectin-1 Contributes to Myocardial Ischemia/Reperfusion Injury by Regulating Macrophage Polarization and Neutrophil Infiltration. Circulation 139(5): 663–678. 10.1161/Circulationaha.118.036044

Fattahi, F., L. M. Frydrych, G. Bian, M. Kalbitz, T. J. Herron, E. A. Malan, M. J. Delano and P. A. Ward (2018). Role of complement C5a and histones in septic cardiomyopathy. Molecular immunology 102: 32–41. 10.1016/j.molimm.2018.06.006

Feng, J., Y. Li, Y. Li, Q. Yin, H. Li, J. Li, B. Zhou, J. Meng, H. Lian, M. Wu, Y. Li, K. Dou, W. Song, B. Lu, L. Liu, S. Hu and Y. Nie (2024). Versican Promotes Cardiomyocyte Proliferation and Cardiac Repair. Circulation 149(13): 1004–1015. 10.1161/CIRCULATIONAHA.123.066298

Gibb, A. A. and B. G. Hill (2018). Metabolic Coordination of Physiological and Pathological Cardiac Remodeling. Circulation Research 123(1): 107–128. 10.1161/Circresaha.118.312017

Hudson, J. E. and E. R. Porrello (2017). Periostin paves the way for neonatal heart regeneration. Cardiovascular Research 113(6): 556–558. 10.1093/cvr/cvx039

Huynh, P., J. D. Hoffmann, T. Gerhardt, M. G. Kiss, F. M. Zuraikat, O. Cohen, C. Wolfram, A. G. Yates, A. Leunig, M. Heiser, L. Gaebel, M. Gianeselli, S. Goswami, A. Khamhoung, J. Downey, S. Yoon, Z. Chen, V. Roudko, T. Dawson, J. Ferreira da Silva, N. J. Ameral, J. Morgenroth-Rebin, D. D’Souza, L. L. Koekkoek, W. Jacob, J. Munitz, D. Lee, J. F. Fullard, M. M. T. van Leent, P. Roussos, S. Kim-Schulze, N. Shah, B. P. Kleinstiver, F. K. Swirski, D. Leistner, M.-P. St-Onge and C. S. McAlpine (2024). Myocardial infarction augments sleep to limit cardiac inflammation and damage. Nature 635(8037): 168–177. 10.1038/s41586-024-08100-w

Jia, D. L., S. Q. Chen, P. Y. Bai, C. T. Luo, J. Liu, A. J. Sun and J. B. Ge (2022). Cardiac Resident Macrophage-Derived Legumain Improves Cardiac Repair by Promoting Clearance and Degradation of Apoptotic Cardiomyocytes After Myocardial Infarction. Circulation 145(20): 1542–1556. 10.1161/Circulationaha.121.057549

Johnson, J., S. Q. K. Law, M. Shojaee, A. S. Hall, S. Bhuiyan, M. B. L. Lim, A. Silva, K. J. W. Kong, M. Schoppet, C. Blyth, H. N. Ranasinghe, N. Sejic, M. J. Chuei, O. C. Tatford, A. Cifuentes-Rius, P. F. James, A. Tester, I. Dixon and G. Lichtfuss (2023). First-in-human clinical trial of allogeneic, platelet-derived extracellular vesicles as a potential therapeutic for delayed wound healing. J Extracell Vesicles 12(7): e12332. 10.1002/jev2.12332

Kanisicak, O., H. Khalil, M. J. Ivey, J. Karch, B. D. Maliken, R. N. Correll, M. J. Brody, S.-C. J Lin, B. J. Aronow, M. D. Tallquist and J. D. Molkentin (2016). Genetic lineage tracing defines myofibroblast origin and function in the injured heart. Nature communications 7: 12260. 10.1038/ncomms12260

Kathiresan, S. and D. Srivastava (2012). Genetics of Human Cardiovascular Disease. Cell 148(6): 1242-1257. 10.1016/j.cell.2012.03.001

Kramann, R. (2022). A map of the human heart after myocardial infarction. Nature. 10.1038/d41586-022-02011-4

Kuppe, C., R. O. Ramirez Flores, Z. Li, S. Hayat, R. T. Levinson, X. Liao, M. T. Hannani, J. Tanevski, F. Wunnemann, J. S. Nagai, M. Halder, D. Schumacher, S. Menzel, G. Schafer, K. Hoeft, M. Cheng, S. Ziegler, X. Zhang, F. Peisker, N. Kaesler, T. Saritas, Y. Xu, A. Kassner, J. Gummert, M. Morshuis, J. Amrute, R. J. A. Veltrop, P. Boor, K. Klingel, L. W. Van Laake, A. Vink, R. M. Hoogenboezem, E. M. J. Bindels, L. Schurgers, S. Sattler, D. Schapiro, R. K. Schneider, K. Lavine, H. Milting, I. G. Costa, J. Saez-Rodriguez and R. Kramann (2022). Spatial multi-omic map of human myocardial infarction. Nature 608(7924): 766–777. 10.1038/s41586-022-05060-x

Li, G., J. Zhang, Z. Zhao, J. Wang, J. Li, W. Xu, Z. Cui, P. Sun, H. Yuan, T. Wang, K. Li, X. Bai, X. Ma, P. Li, Y. Fu, Y. Cao, H. Bao, D. Li, Z. Liu, N. Zhu, L. Tang and Z. Lu (2024). RNF144B negatively regulates antiviral immunity by targeting MDA5 for autophagic degradation. EMBO reports 25(10): 4594–4624. 10.1038/s44319-024-00256-w

Li, W., X. Lou, Y. Zha, Y. Qin, J. Zha, L. Hong, Z. Xie, S. Yang, C. Wang, J. An, Z. Zhang and S. Qiao (2023). Single-cell RNA-seq of heart reveals intercellular communication drivers of myocardial fibrosis in diabetic cardiomyopathy. eLife 12. 10.7554/eLife.80479

Liu, A., Z. Chen, X. Li, C. Xie, Y. Chen, X. Su, Y. Chen, M. Zhang, J. Chen, T. Yang, J. Shen and H. Huang (2023). C5a-C5aR1 induces endoplasmic reticulum stress to accelerate vascular calcification via PERK-eIF2alpha-ATF4-CREB3L1 pathway. Cardiovascular research 119(15): 2563–2578. 10.1093/cvr/cvad133

Martini, E. (2023). A machine learning-based system to improve myocardial infarction diagnosis. Nature cardiovascular research 2(6): 490. 10.1038/s44161-023-00291-1

McCarroll, C. S., W. He, K. Foote, A. Bradley, K. McGlynn, F. Vidler, C. Nixon, K. Nather, C. Fattah, A. Riddell, P. Bowman, E. B. Elliott, M. Bell, C. Hawksby, S. M. MacKenzie, L. J. Morrison, A. Terry, K. Blyth, G. L. Smith, M. W. McBride, T. Kubin, T. Braun, S. A. Nicklin, E. R. Cameron and C. M. Loughrey (2018). Runx1 Deficiency Protects Against Adverse Cardiac Remodeling After Myocardial Infarction. Circulation 137(1): 57–70. 10.1161/CIRCULATIONAHA.117.028911

McCarthy, C. P., M. Vaduganathan and J. L. Januzzi, Jr. (2018). Type 2 Myocardial Infarction-Diagnosis, Prognosis, and Treatment. Jama 320(5): 433–434. 10.1001/jama.2018.7125

Morange, P. E., N. Saut, M. C. Alessi, J. S. Yudkin, M. Margaglione, G. Di Minno, A. Hamsten, S. E. Humphries, D. A. Tregouet and I. Juhan-Vague (2007). Association of plasminogen activator inhibitor (PAI)-1 (SERPINE1) SNPs with myocardial infarction, plasma PAI-1, and metabolic parameters: the HIFMECH study. Arteriosclerosis, thrombosis, and vascular biology 27(10): 2250–2257. 10.1161/ATVBAHA.107.149468

Natarajan, N., Y. Abbas, D. M. Bryant, J. M. Gonzalez-Rosa, M. Sharpe, A. Uygur, L. H. Cocco-Delgado, N. N. Ho, N. P. Gerard, C. J. Gerard, C. A. MacRae, C. E. Burns, C. G. Burns, J. L. Whited and R. T. Lee (2018). Complement Receptor C5aR1 Plays an Evolutionarily Conserved Role in Successful Cardiac Regeneration. Circulation 137(20): 2152–2165. 10.1161/CIRCULATIONAHA.117.030801

Nguyen, N. U. N., D. C. Canseco, F. Xiao, Y. Nakada, S. J. Li, N. T. Lam, S. A. Muralidhar, J. J. Savla, J. A. Hill, V. T. Le, K. A. Zidan, H. W. El-Feky, Z. N. Wang, M. S. Ahmed, M. E. Hubbi, I. Menendez-Montes, J. Moon, S. R. Ali, V. T. Le, E. Villalobos, M. S. Mohamed, W. M. Elhelaly, S. Thet, C. G. Anene-Nzelu, W. L. W. Tan, R. S. Foo, X. Meng, M. Kanchwala, C. Xing, J. Roy, M. S. Cyert, B. A. Rothermel and H. A. Sadek (2020). A calcineurin-Hoxb13 axis regulates growth mode of mammalian cardiomyocytes. Nature 582(7811): 271-+. 10.1038/s41586-020-2228-6

Niyonzima, N., J. Rahman, N. Kunz, E. E. West, T. Freiwald, J. Desai, N. S. Merle, A. Gidon, B. Sporsheim, M. S. Lionakis, K. Evensen, B. Lindberg, K. Skagen, M. Skjelland, P. Singh, M. Haug, M. M. Ruseva, M. Kolev, J. Bibby, O. Marshall, B. O’Brien, N. Deeks, B. Afzali, R. J. Clark, T. M. Woodruff, M. Pryor, Z. H. Yang, A. T. Remaley, T. E. Mollnes, S. M. Hewitt, B. Y. Yan, M. Kazemian, M. G. Kiss, C. J. Binder, B. Halvorsen, T. Espevik and C. Kemper (2021). Mitochondrial C5aR1 activity in macrophages controls IL-1β production underlying sterile inflammation. Science Immunology 6(66). 10.1126/sciimmunol.abf2489

Nording, H., L. Baron, D. Haberthur, F. Emschermann, M. Mezger, M. Sauter, R. Sauter, J. Patzelt, K. Knoepp, A. Nording, M. Meusel, R. Meyer-Saraei, R. Hlushchuk, D. Sedding, O. Borst, I. Eitel, C. M. Karsten, R. Feil, B. Pichler, J. Erdmann, A. Verschoor, E. Chavakis, T. Chavakis, P. von Hundelshausen, J. Kohl, M. Gawaz and H. F. Langer (2021). The C5a/C5a receptor 1 axis controls tissue neovascularization through CXCL4 release from platelets. Nature communications 12(1): 3352. 10.1038/s41467-021-23499-w

Peet, C., A. Ivetic, D. I. Bromage and A. M. Shah (2020). Cardiac monocytes and macrophages after myocardial infarction. Cardiovascular Research 116(6): 1101–1112. 10.1093/cvr/cvz336

Prabhu, S. D. and N. G. Frangogiannis (2016). The Biological Basis for Cardiac Repair After Myocardial Infarction From Inflammation to Fibrosis. Circulation Research 119(1): 91–112. 10.1161/Circresaha.116.303577

Pu, Z., X. Bao, S. Xia, P. Shao and Y. Xu (2022). Serpine1 Regulates Peripheral Neutrophil Recruitment and Acts as Potential Target in Ischemic Stroke. Journal of inflammation research 15: 2649–2663. 10.2147/JIR.S361072

Qian, L. (2016). Cellular reprogramming improves cardiac function in a mouse model of myocardial infarction. Science 352(6292): 1400–1401. 10.1126/science.aag1213

Reboll, M. R., S. Klede, M. H. Taft, C. L. Cai, L. J. Field, K. J. Lavine, A. L. Koenig, J. Fleischauer, J. Meyer, A. Schambach, H. W. Niessen, M. Kosanke, J. van den Heuvel, A. Pich, J. Bauersachs, X. K. Wu, L. Q. Zheng, Y. Wang, M. Korf-Klingebiel, F. Polten and K. C. Wollert (2022). Meteorin-like promotes heart repair through endothelial KIT receptor tyrosine kinase. Science 376(6599): 1343-+. 10.1126/science.abn3027

Riddell, A., M. McBride, T. Braun, S. A. Nicklin, E. Cameron, C. M. Loughrey and T. P. Martin (2020). RUNX1: an emerging therapeutic target for cardiovascular disease. Cardiovascular research 116(8): 1410–1423. 10.1093/cvr/cvaa034

Ruiz-Villalba, A., J. P. Romero, S. C. Hernandez, A. Vilas-Zornoza, N. Fortelny, L. Castro-Labrador, P. San Martin-Uriz, E. Lorenzo-Vivas, P. Garcia-Olloqui, M. Palacio, J. J. Gavira, G. Bastarrika, S. Janssens, M. Wu, E. Iglesias, G. Abizanda, X. M. de Morentin, M. Lasaga, N. Planell, C. Bock, D. Alignani, G. Medal, I. Prudovsky, Y.-R. Jin, S. Ryzhov, H. Yin, B. Pelacho, D. Gomez-Cabrero, V. Lindner, D. Lara-Astiaso and F. Prosper (2020). Single-Cell RNA Sequencing Analysis Reveals a Crucial Role for CTHRC1 (Collagen Triple Helix Repeat Containing 1) Cardiac Fibroblasts After Myocardial Infarction. Circulation 142(19): 1831–1847. 10.1161/CIRCULATIONAHA.119.044557

Salari, N., F. Morddarvanjoghi, A. Abdolmaleki, S. Rasoulpoor, A. A. Khaleghi, L. A. Hezarkhani, S. Shohaimi and M. Mohammadi (2023). The global prevalence of myocardial infarction: a systematic review and meta-analysis. BMC Cardiovasc Disord 23(1): 206. 10.1186/s12872-023-03231-w

Schlotter, F., K. Huber, C. Hassager, S. Halvorsen, P. Vranckx, J. Poss, K. Krychtiuk, R. Lorusso, N. Bonaros, P. A. Calvert, M. Montorfano and H. Thiele (2024). Ventricular septal defect complicating acute myocardial infarction: diagnosis and management. A Clinical Consensus Statement of the Association for Acute CardioVascular Care (ACVC) of the ESC, the European Association of Percutaneous Cardiovascular Interventions (EAPCI) of the ESC and the ESC Working Group on Cardiovascular Surgery. European heart journal 45(28): 2478–2492. 10.1093/eurheartj/ehae363

Vergallo, R. and C. Patrono (2023). Machine learning and myocardial infarction diagnosis: sometimes you can’t make it on your own. European heart journal 44(35): 3309–3310. 10.1093/eurheartj/ehad467

Virani (2023). 2023 AHA/ACC/ACCP/ASPC/NLA/PCNA Guideline for the Management of Patients With Chronic Coronary Disease: A Report of the American Heart Association/American College of Cardiology Joint Committee on Clinical Practice Guidelines (vol 148, pg e9, 2023). Circulation 148(13): E148-E148. 10.1161/Cir.0000000000001183

Wang, H. and L. Dou (2024). Single-cell RNA sequencing reveals hub genes of myocardial infarction-associated endothelial cells. BMC cardiovascular disorders 24(1): 70. 10.1186/s12872-024-03727-z

Xu, Y., K. Jiang, F. Su, R. Deng, Z. Cheng, D. Wang, Y. Yu and Y. Xiang (2023). A transient wave of Bhlhe41(+) resident macrophages enables remodeling of the developing infarcted myocardium. Cell Rep 42(10): 113174. 10.1016/j.celrep.2023.113174

Yokota, T., J. McCourt, F. Ma, S. Ren, S. Li, T.-H. Kim, Y. Z. Kurmangaliyev, R. Nasiri, S. Ahadian, T. Nguyen, X. H. M. Tan, Y. Zhou, R. Wu, A. Rodriguez, W. Cohn, Y. Wang, J. Whitelegge, S. Ryazantsev, A. Khademhosseini, M. A. Teitell, P.-Y. Chiou, D. E. Birk, A. C. Rowat, R. H. Crosbie, M. Pellegrini, M. Seldin, A. J. Lusis and A. Deb (2020). Type V Collagen in Scar Tissue Regulates the Size of Scar after Heart Injury. Cell 182(3): 545–562.e523. 10.1016/j.cell.2020.06.030

Yokota, T., J. McCourt, F. Y. Ma, S. X. Ren, S. Li, T. H. Kim, Y. Z. Kurmangaliyev, R. Nasiri, S. Ahadian, T. Nguyen, X. H. M. Tan, Y. G. Zhou, R. M. Wu, A. Rodriguez, W. Cohn, Y. B. Wang, J. Whitelegge, S. Ryazantsev, A. Khademhosseini, M. A. Teitell, P. Y. Chiou, D. E. Birk, A. C. Rowat, R. H. Crosbie, M. Pellegrini, M. Seldin, A. J. Lusis and A. Deb (2020). Type V Collagen in Scar Tissue Regulates the Size of Scar after Heart Injury. Cell 182(3): 545-+. 10.1016/j.cell.2020.06.030

Zhao, X. Y., T. Williamson, Y. Q. Gong, J. A. Epstein and Y. Fan (2024). Immunomodulatory Therapy for Ischemic Heart Disease. Circulation 150(13): 1050–1058. 10.1161/Circulationaha.124.070368

Zhuang, H., Z. Zhang, W. Wang and H. Qu (2024). RNF144B-mediated p21 degradation regulated by HDAC3 contribute to enhancing ovarian cancer growth and metastasis. Tissue & cell 86: 102277. 10.1016/j.tice.2023.102277

