## supplemental for "Single-cell transcriptomics and machine learning reveal RNF144B and C5AR1 as immune-related biomarkers and therapeutic targets in myocardial infarction": Supplementary fig.pdf

Supplemental fig.1

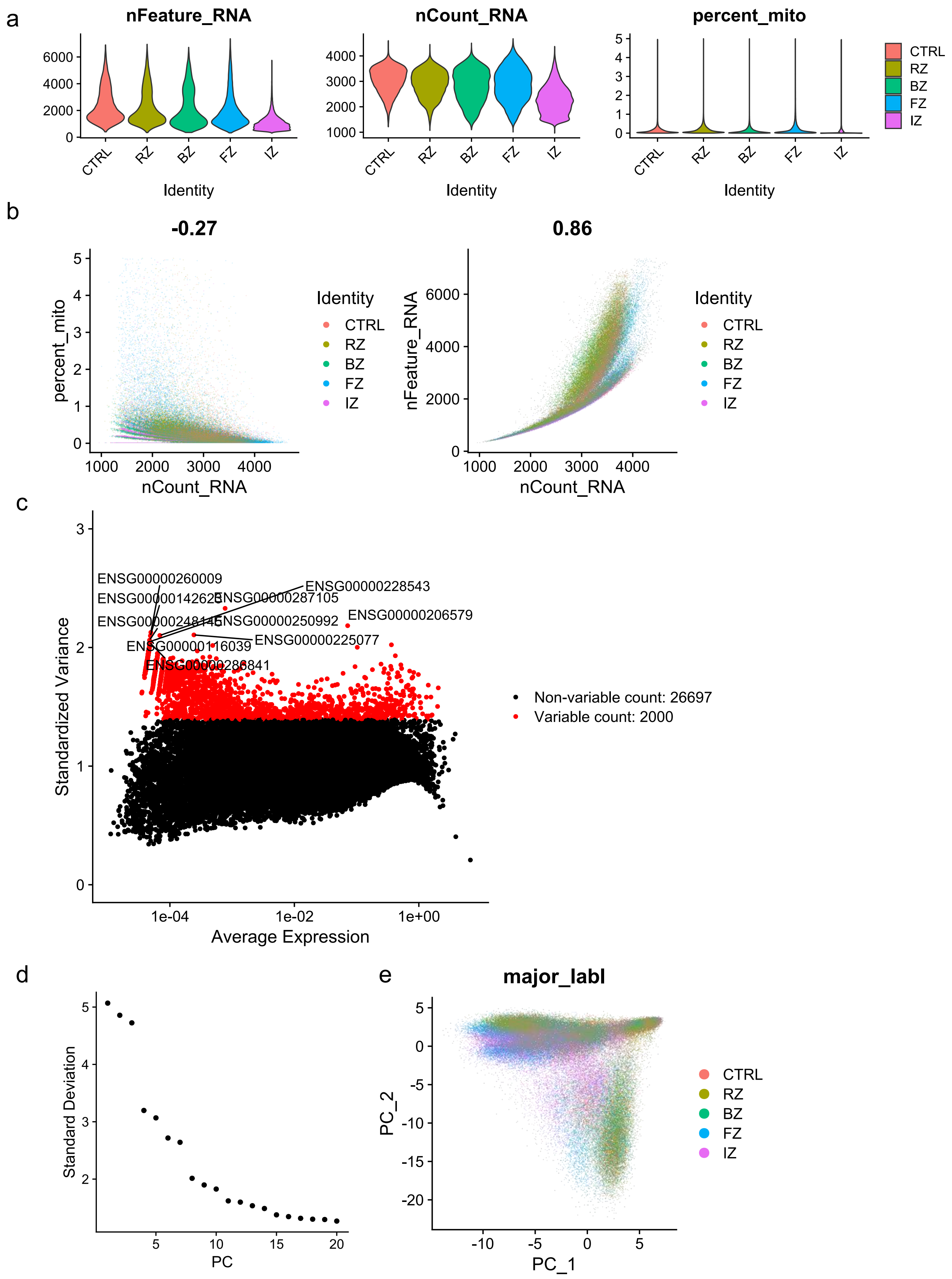

Supplemental fig.2

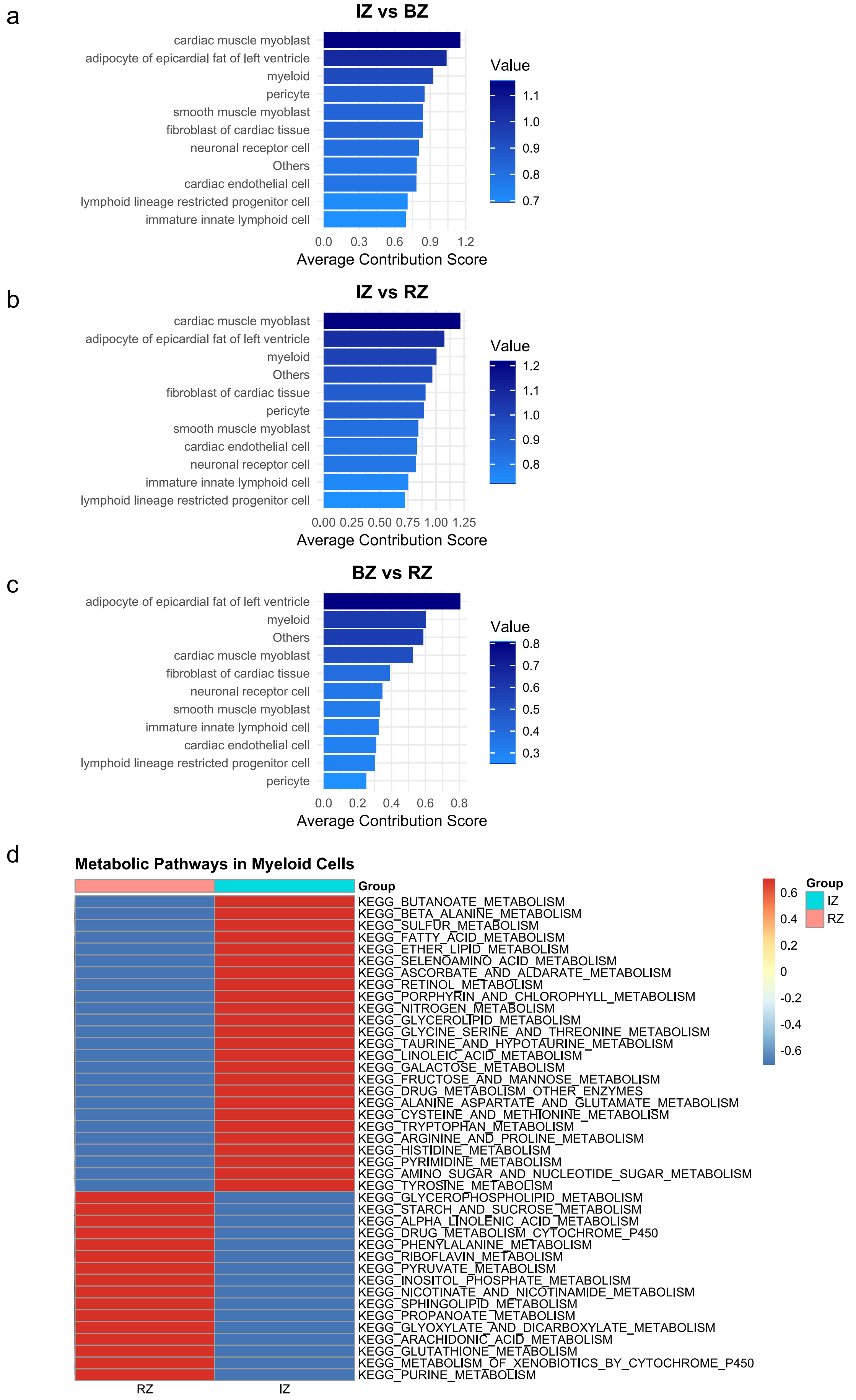

Supplemental fig.3

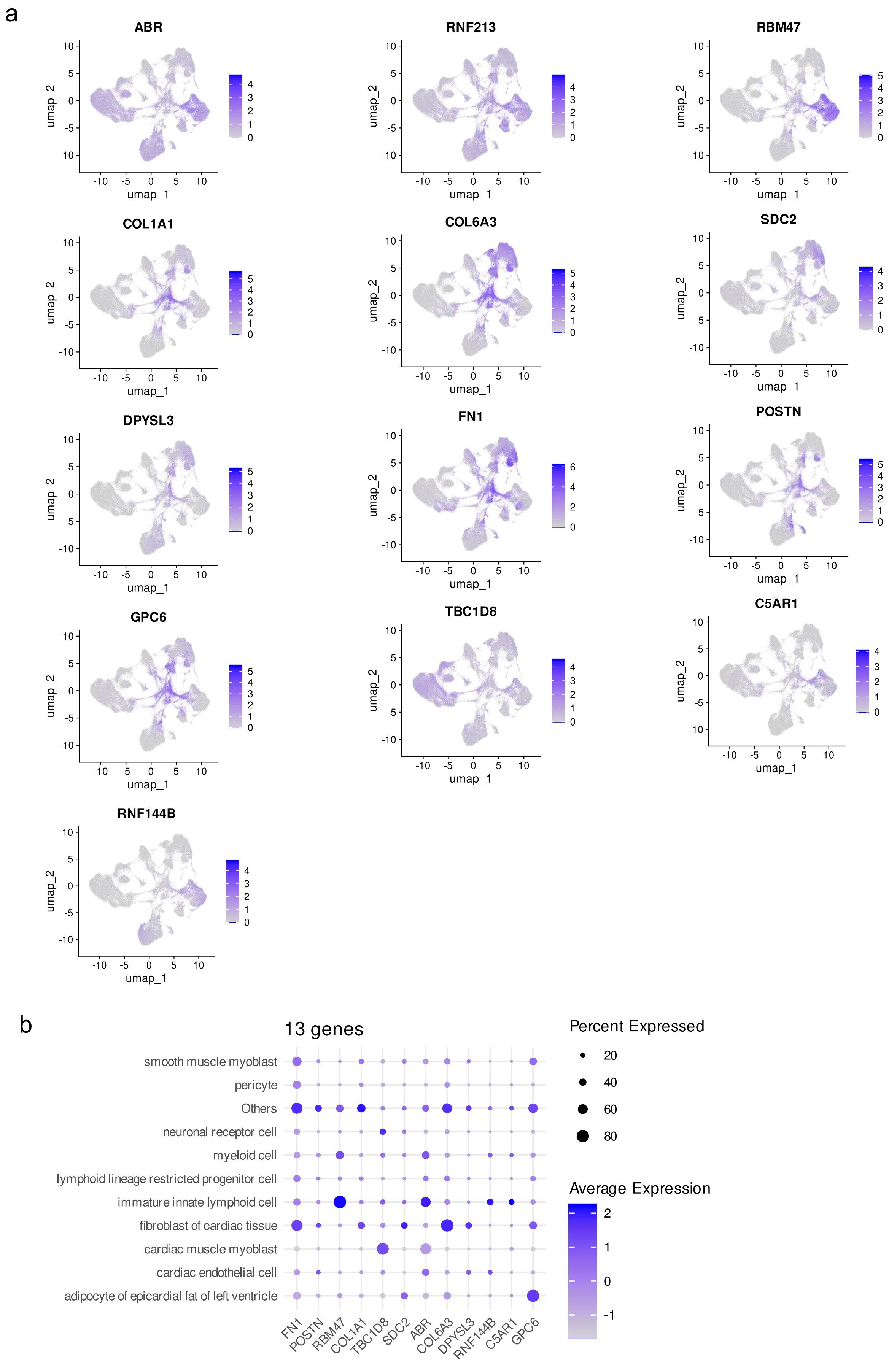

### Supplemental fig.4

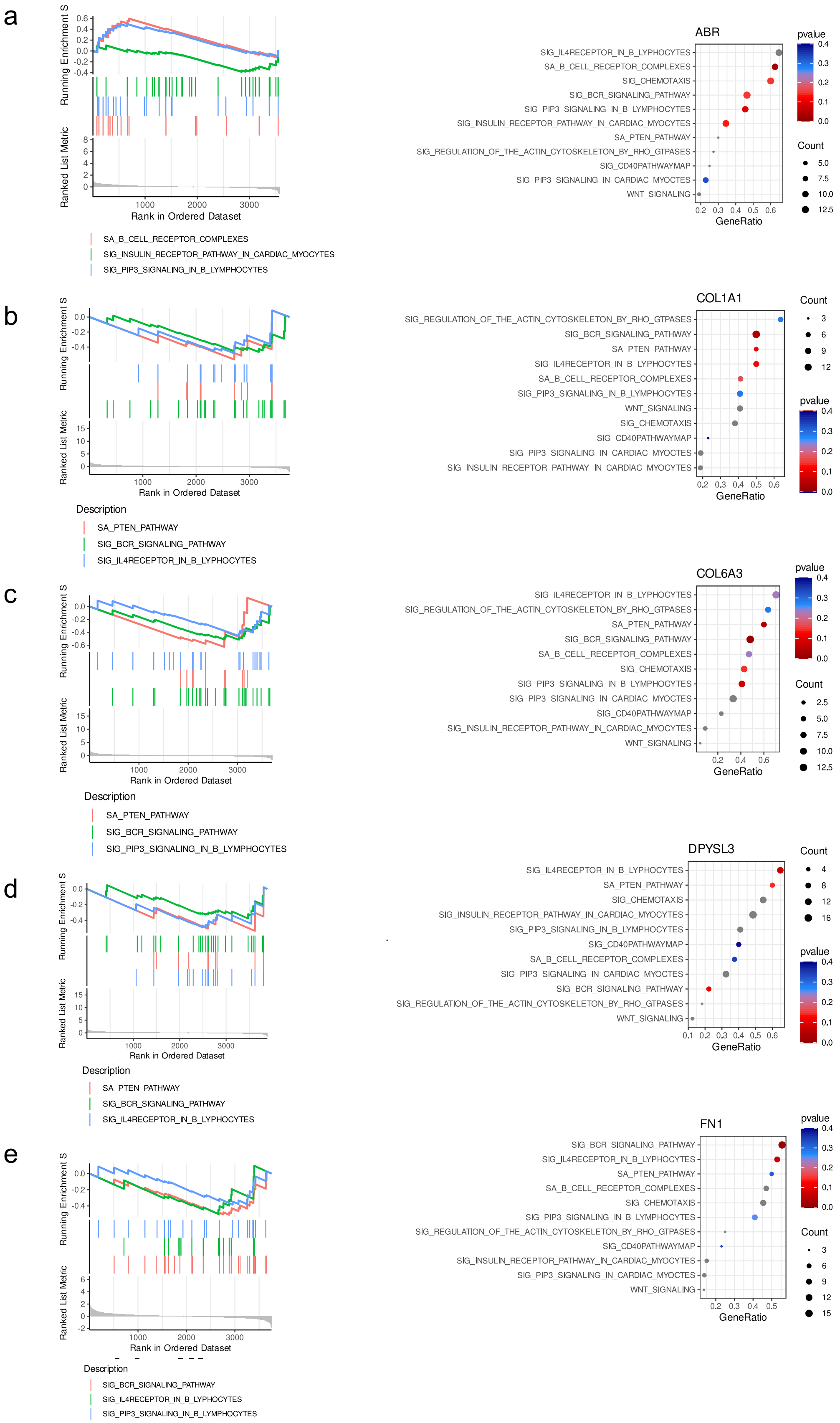

### Supplemental fig.4

f

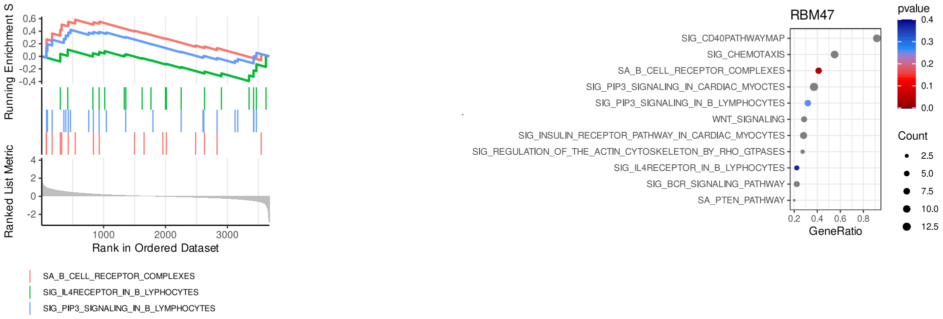


g

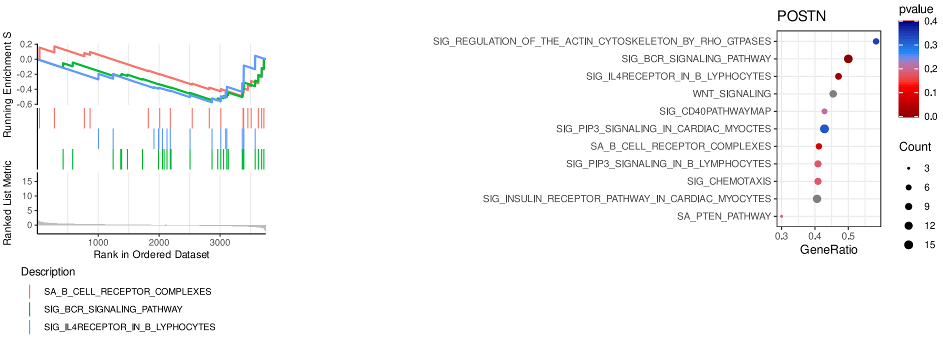


h

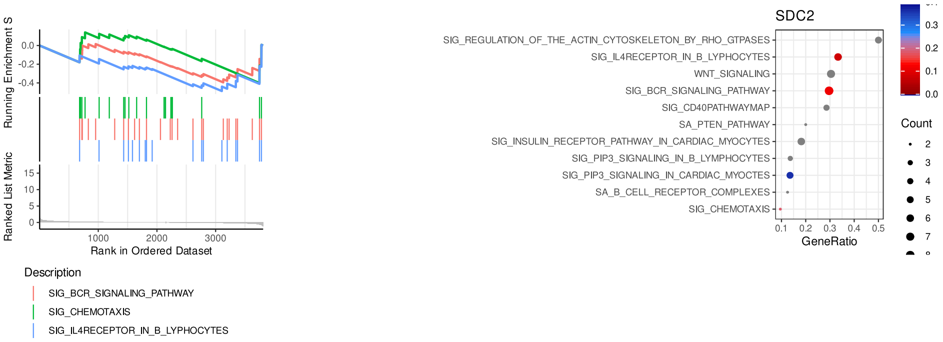


i

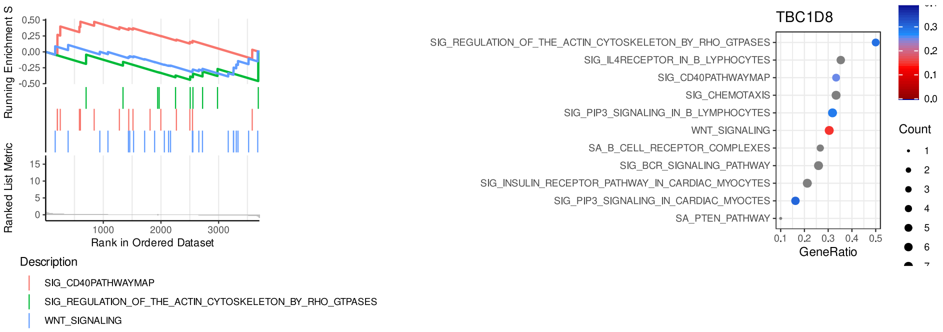


j

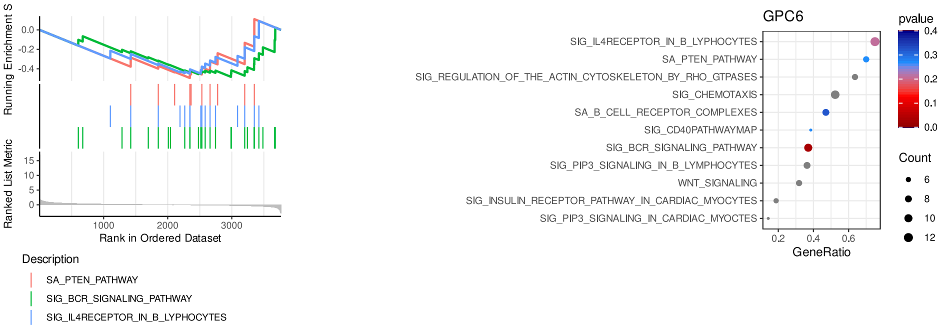

Supplemental fig.5

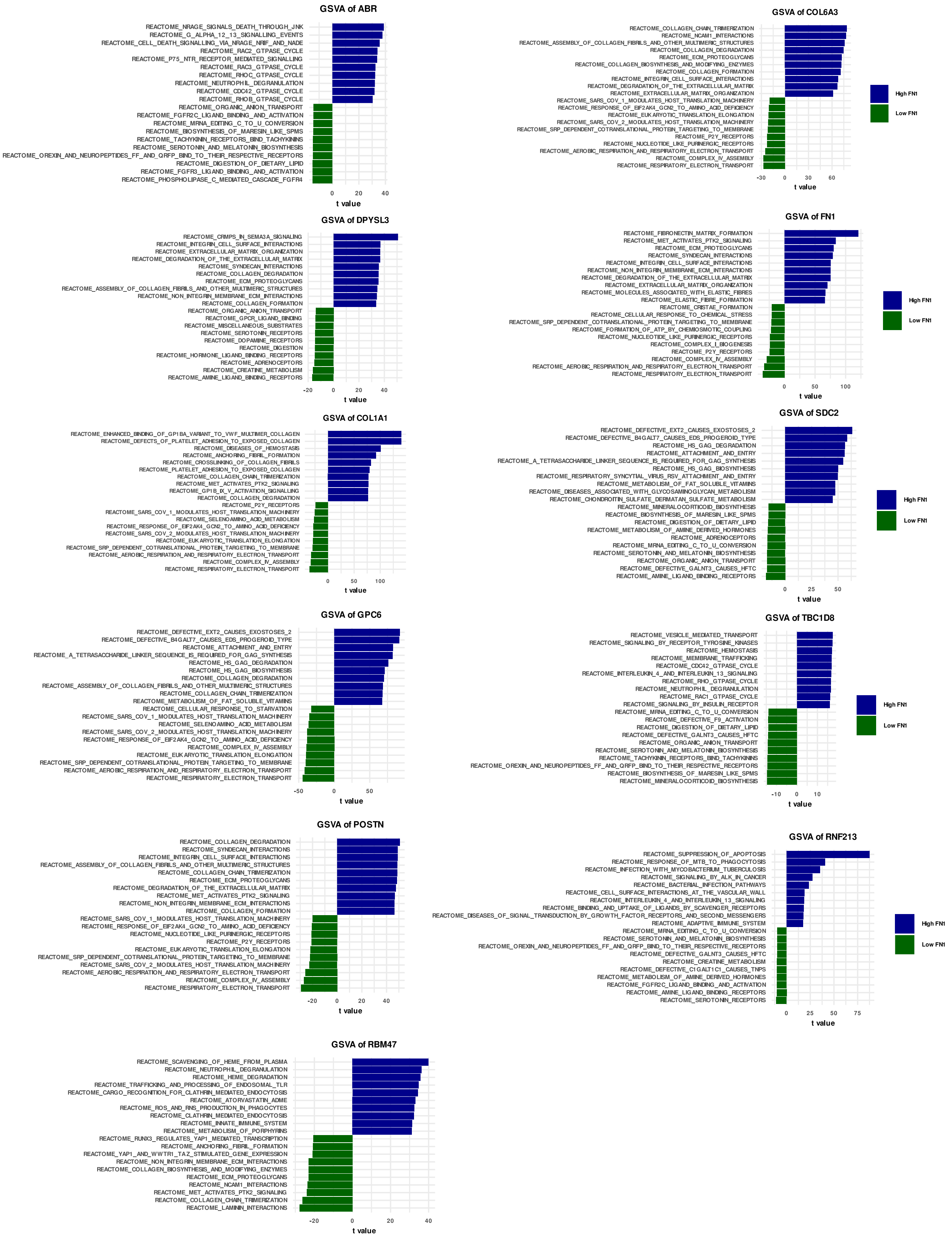

Supplemental fig.6

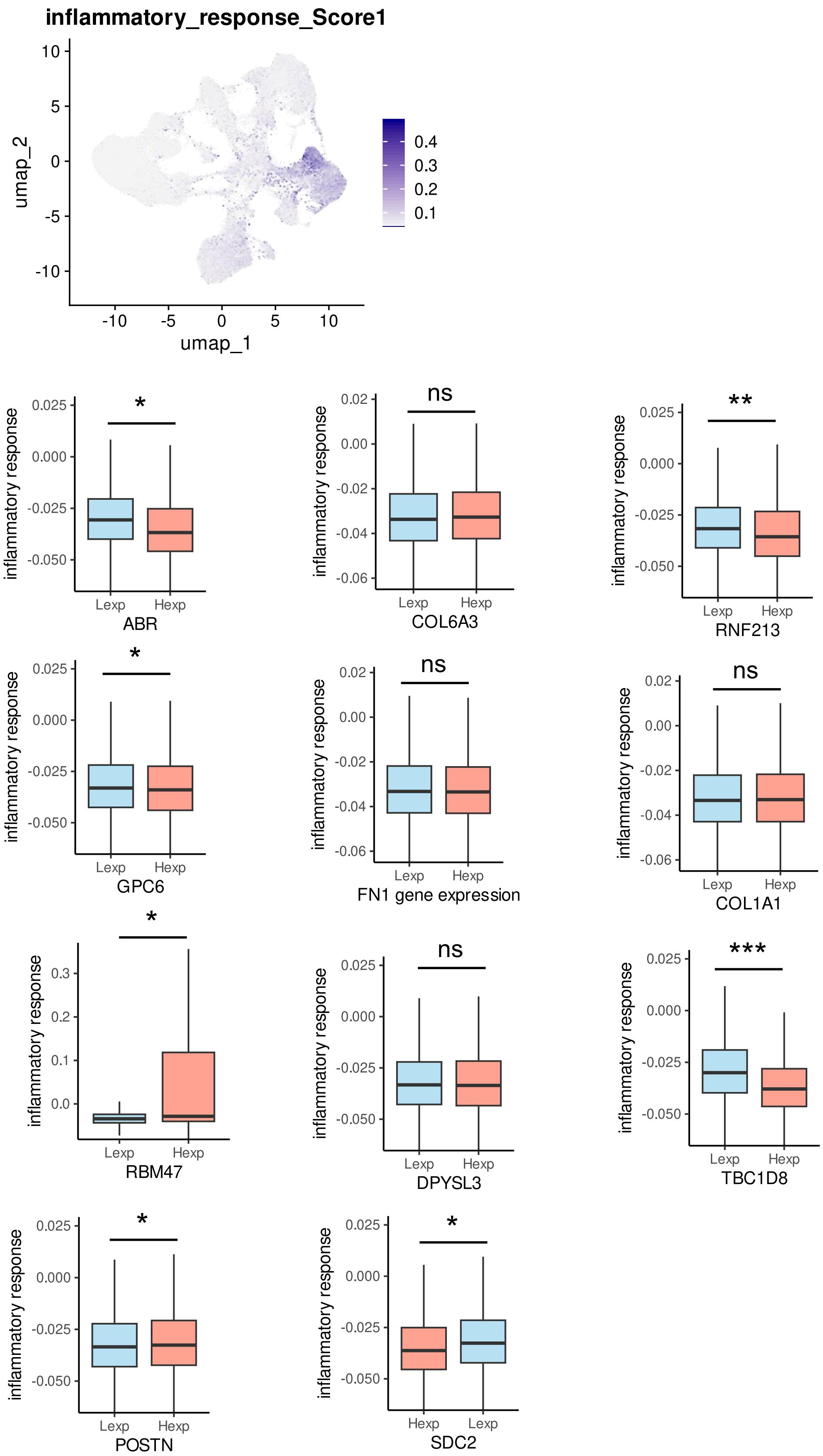

Supplemental fig.7

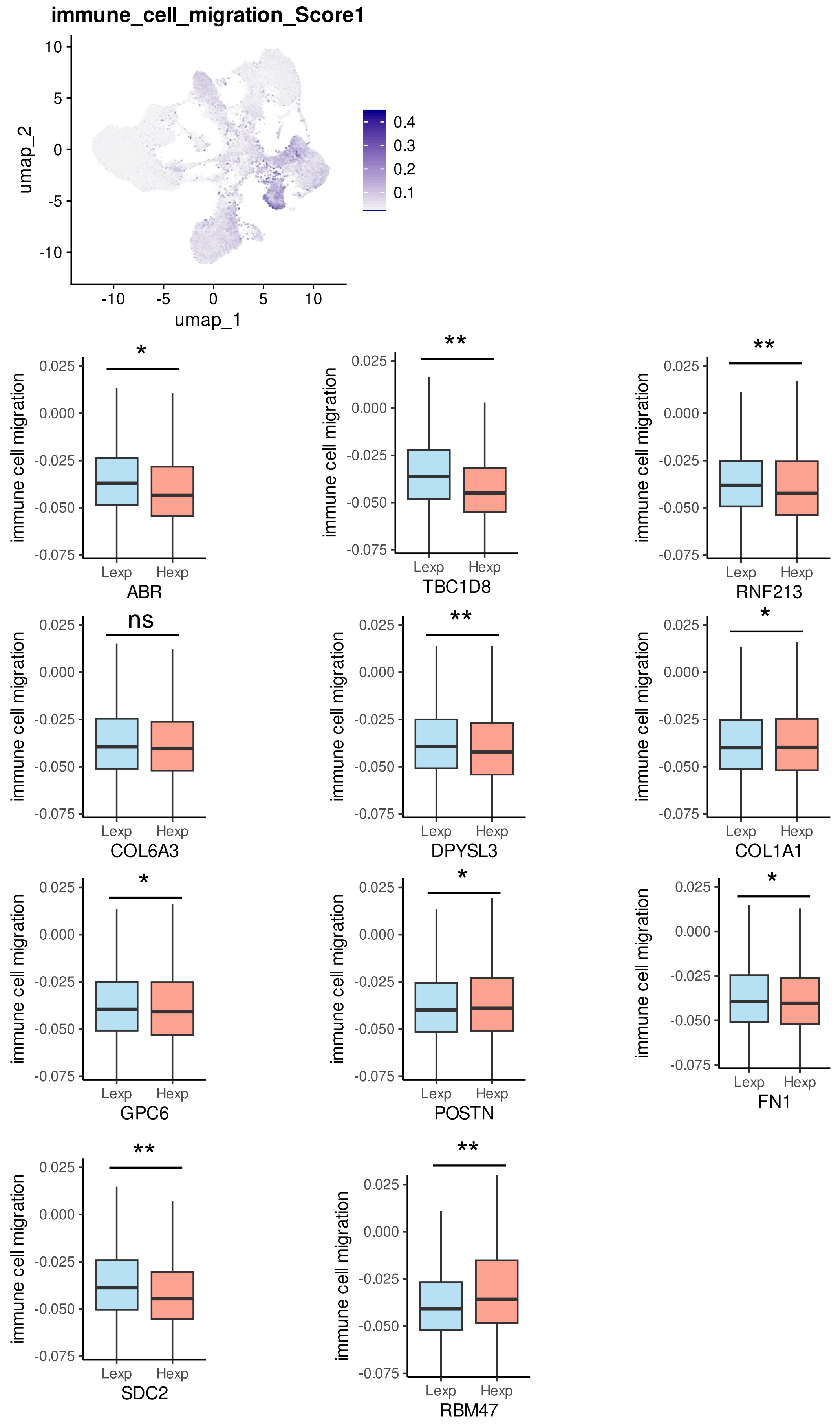

Supplemental fig.8

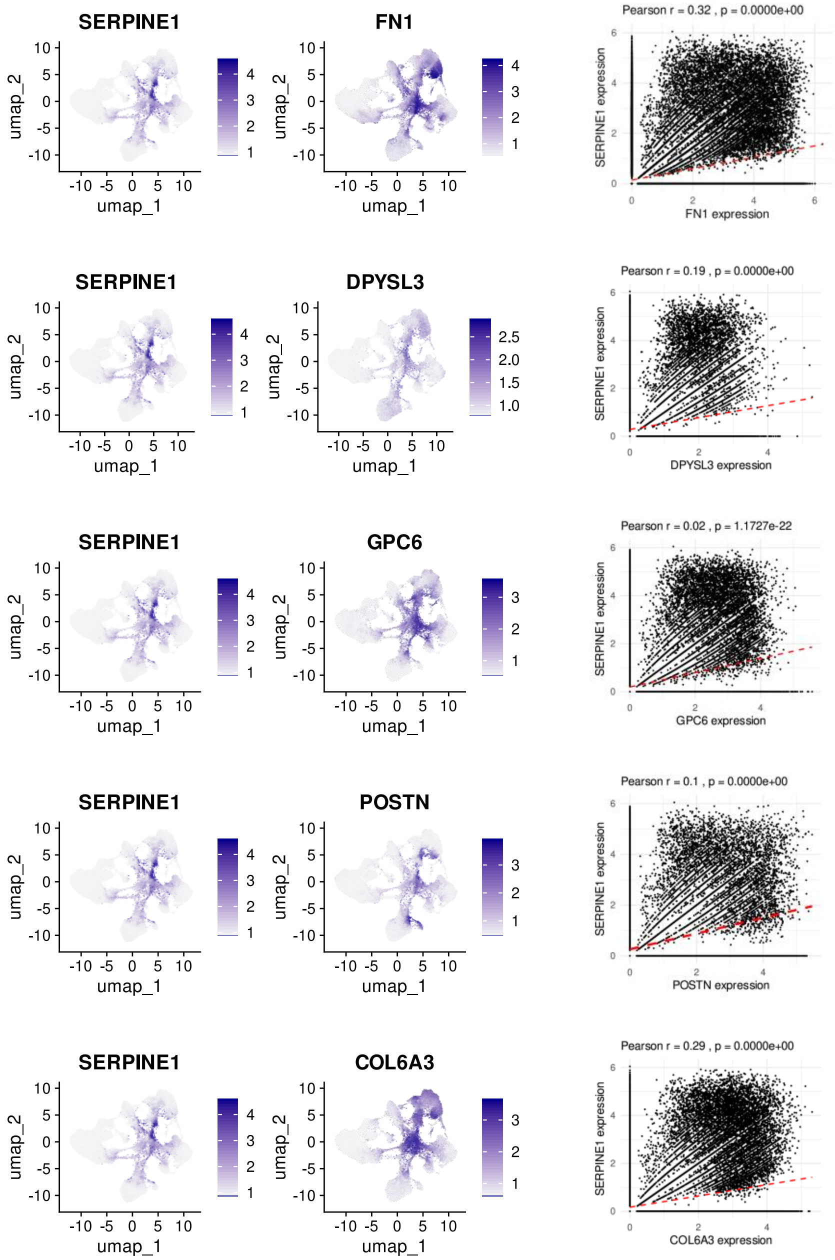

Supplemental fig.8

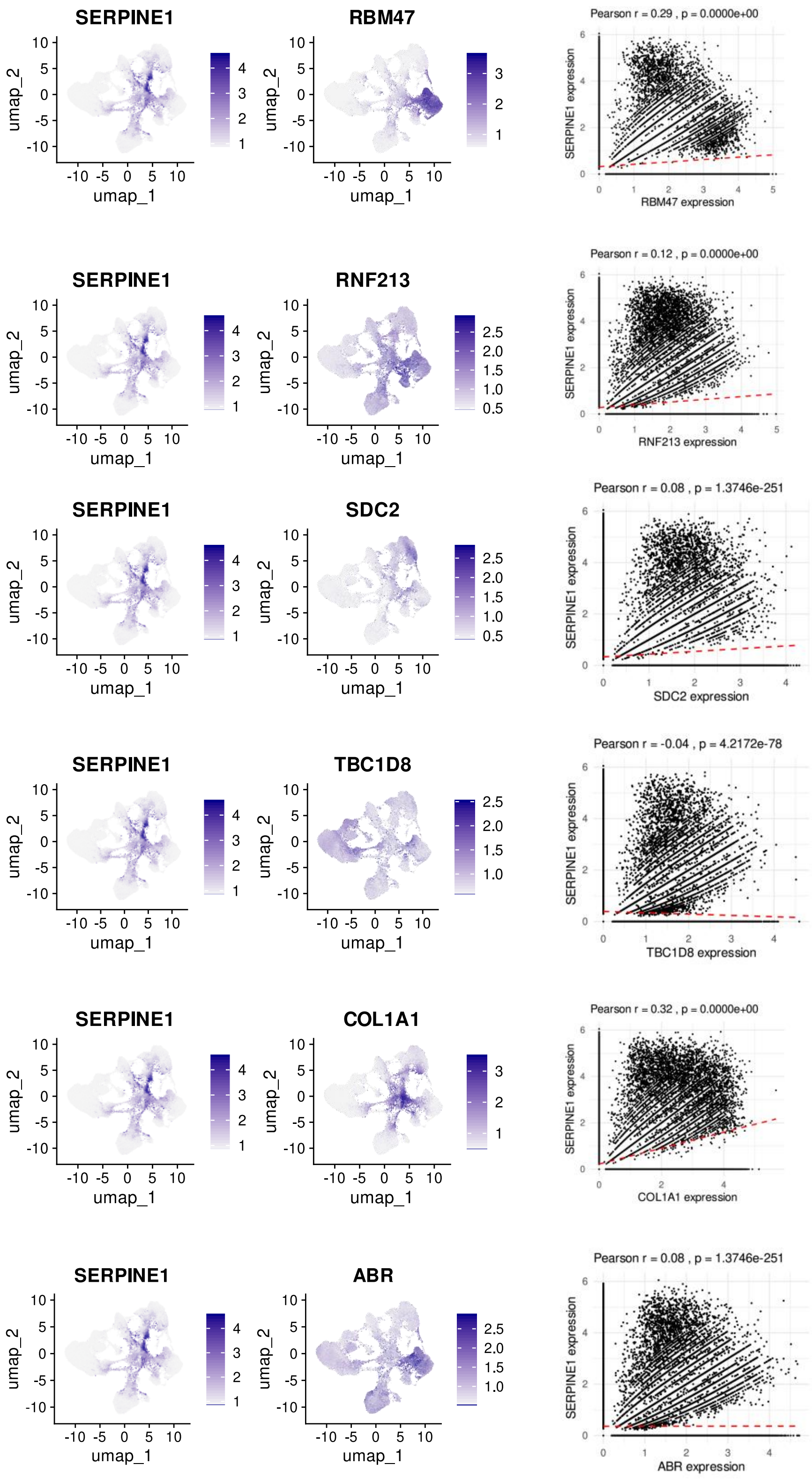

Supplemental fig.9

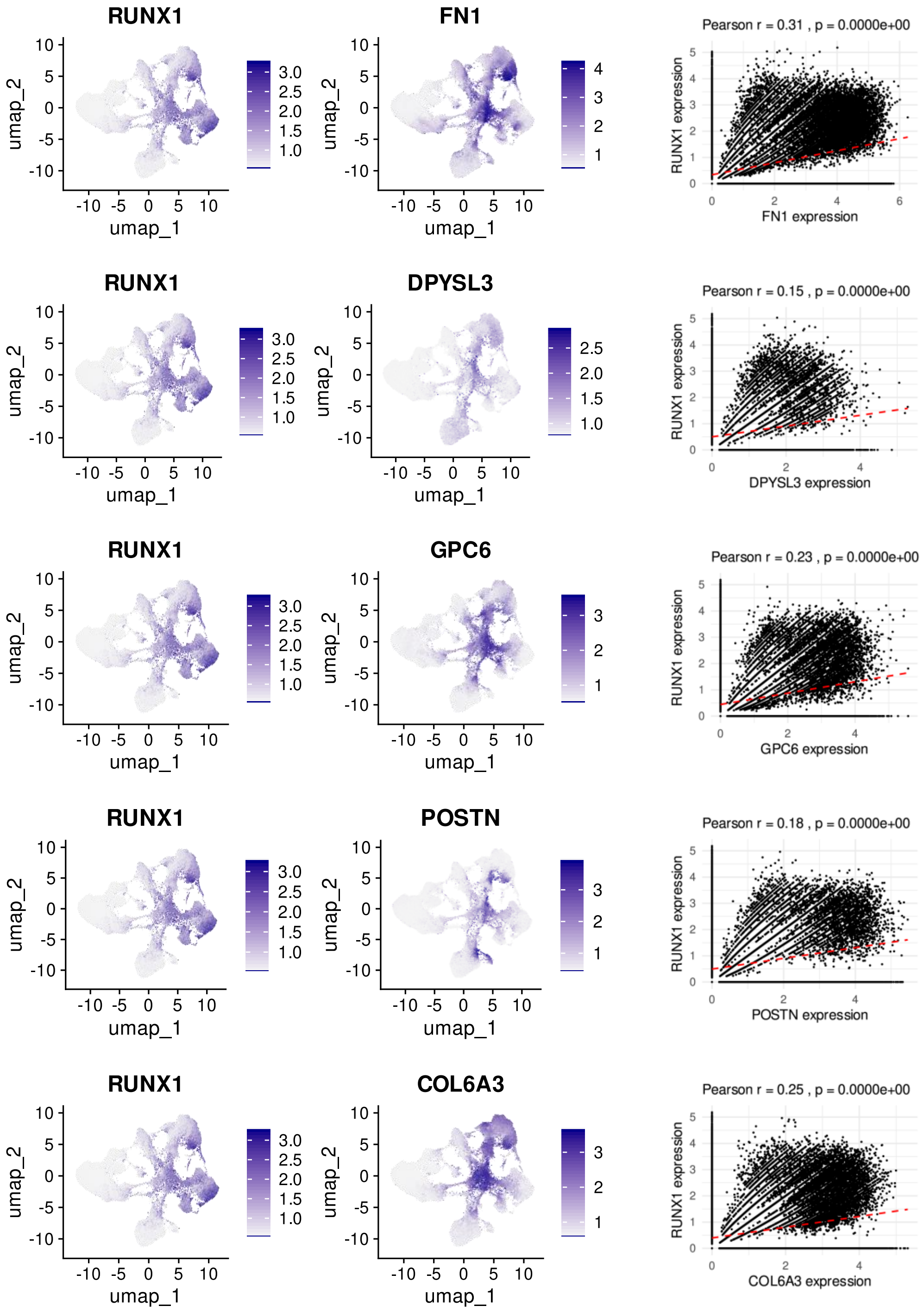

Supplemental fig.9

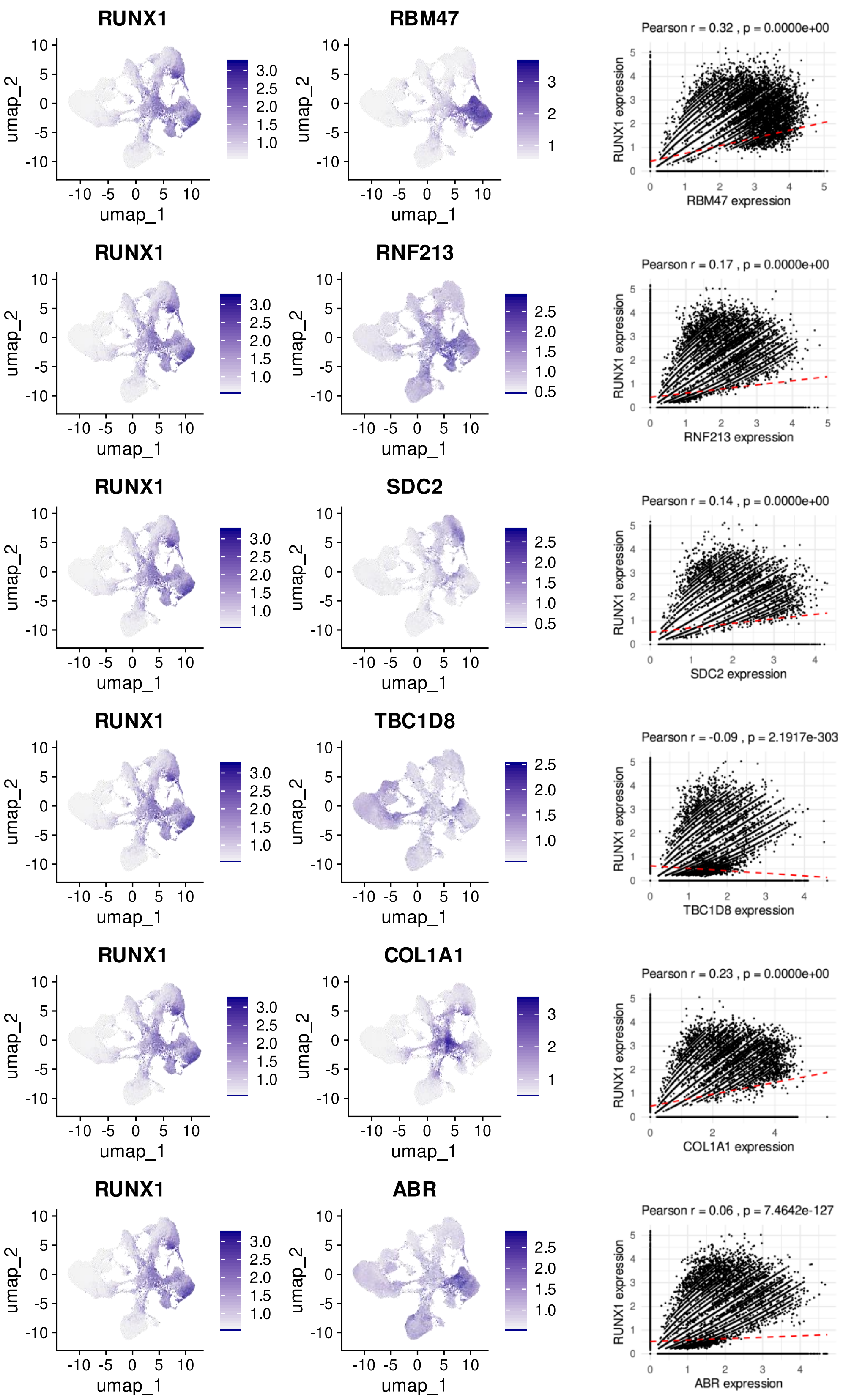
